# Light-powered reactivation of flagella and contraction of microtubules network: towards building an artificial cell

**DOI:** 10.1101/2020.07.20.212191

**Authors:** Raheel Ahmad, Christin Kleineberg, Vahid Nasirimarekani, Yu-Jung Su, Samira Goli Pozveh, Albert Bae, Kai Sundmacher, Eberhard Bodenschatz, Isabella Guido, Tanja Vidakovic-Koch, Azam Gholami

## Abstract

Artificial systems capable of self-sustained movement with self-sufficient energy are of high interest with respect to the development of many challenging applications including medical treatments but also technical applications. The bottom-up assembly of such systems in the context of synthetic biology is still a challenging task. In this work, we demonstrate the biocompatibility and efficiency of an artificial light-driven energy module and a motility functional unit by integrating light-switchable photosynthetic vesicles with demembranated flagella that provide ATP for dynein molecular motors upon illumination. The flagellar propulsion is coupled to the beating frequency and dynamic ATP synthesis in response to illumination allows us to control beating frequency of flagella in a light-dependent manner. In addition, we verified the functionality of light-powered synthetic vesicles in *in vitro* motility assays by encapsulating microtubules assembled with force-generating kinesin-1 motors and the energy module to investigate the dynamics of a contractile filamentous network in cell-like compartments by optical stimulation. Integration of this photosynthetic system with various biological building blocks such as cytoskeletal filaments and molecular motors may contribute to the bottom-up synthesis of artificial cells that are able to undergo motor-driven morphological deformations and exhibit directional motion in a light-controllable fashion.

**Graphical TOC Entry:** 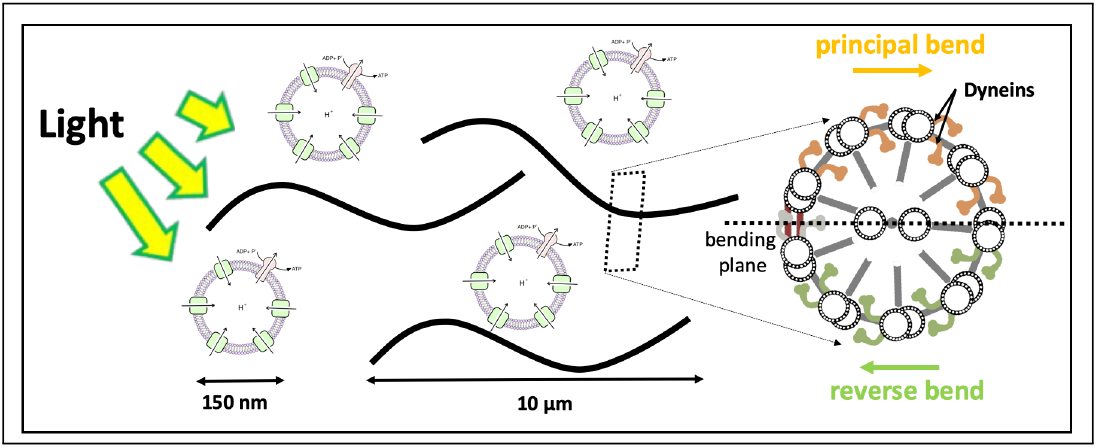

## Introduction

Bottom-up synthetic biology is an emerging field of science that focuses on the engineering of life-like structures. These artificial constructs mimic some of the essential life processes such as reproduction, growth, motility, etc. The reconstitution of each of these processes is quite complex, so the bottom-up synthetic biology initially focuses on sub-systems (functional modules), which are hand-tailored to partially or entirely mimic some of the essential life processes^1–4^. Later on, these functional modules can be combined to build higher hierarchical structures. Artificial systems capable of moving forward are highly attractive for different applications. Examples are smart drug delivery vehicles or so-called designer immune cells which would be capable of detecting and moving towards a chemical signal or tumor^5^. In these applications motility is only one of the functionalities, but even its reconstitution is quite challenging since it embraces, not only the assembly of molecular motors (a motility functional module) which enable movement, but also the energy supply (energy functional module).

In nature, different motility mechanisms have evolved, including polarized assembly and disassembly of bio-polymers for directional motion of amoeboid cells^6,7^, or propulsion powered by tiny hair-like organelles cilia and flagella which perform whip-like motion to provide motility^8^. Arrays of ciliated cells work together to generate directed fluid transport. This includes removal of pollutants in the trachea^9^ or the shuttling and delivery of messenger-containing cerebral spinal fluid (CSF) in our brains^10^. Besides the fluid transport, the regular beating pattern of cilia and flagella propels microorganisms such as spermatozoa, or unicellular organism such as *Paramecium* and green algae *Chlamydomonas reinhardtii*^11–14^. Cilia and flagella are highly conserved organelles composed of a microtubule-based structure called axoneme which is covered by a plasma membrane^19,20^. Nine microtubule doublets, each formed of a complete A-microtubule and an incomplete B-microtubule, are cylindrically arranged around a central pair of singlet microtubules (Figure 1). The diameter of an axoneme is around 200 nm, and the adjacent microtubule doublets are spaced 30 nm away from each other21. The functionality and structural properties of cilia and flagella are the same and can be classified as motile (9+2) and immotile sensory (9+0) cilia. To date, more than 250 constituent proteins have been found to contribute to the highly ordered and precisely assembled structure of axoneme^22^. The microtubule doublets and the dynein molecular motors are the most important components of axonemes^23,24^. Axonemal dyneins are highly specific for ATP as a substrate^25,26^. In the presence of ATP, dynein molecular motors, which are statically bound to the A tubules of each doublet, anchor transiently to the B tubules of the neighboring doublet and generate internal stresses to displace adjacent microtubules relative to each other. However, since microtubule doublets are mechanically connected to the neighboring doublets and cannot slide freely^27^, this sliding force is converted into a bending motion (Figure 1C). The bottom-up reconstitution of flagella is of significant complexity, so we instead isolated this functional module from green algae *Chlamydomonas reinhardtii.*

**Figure 1:**
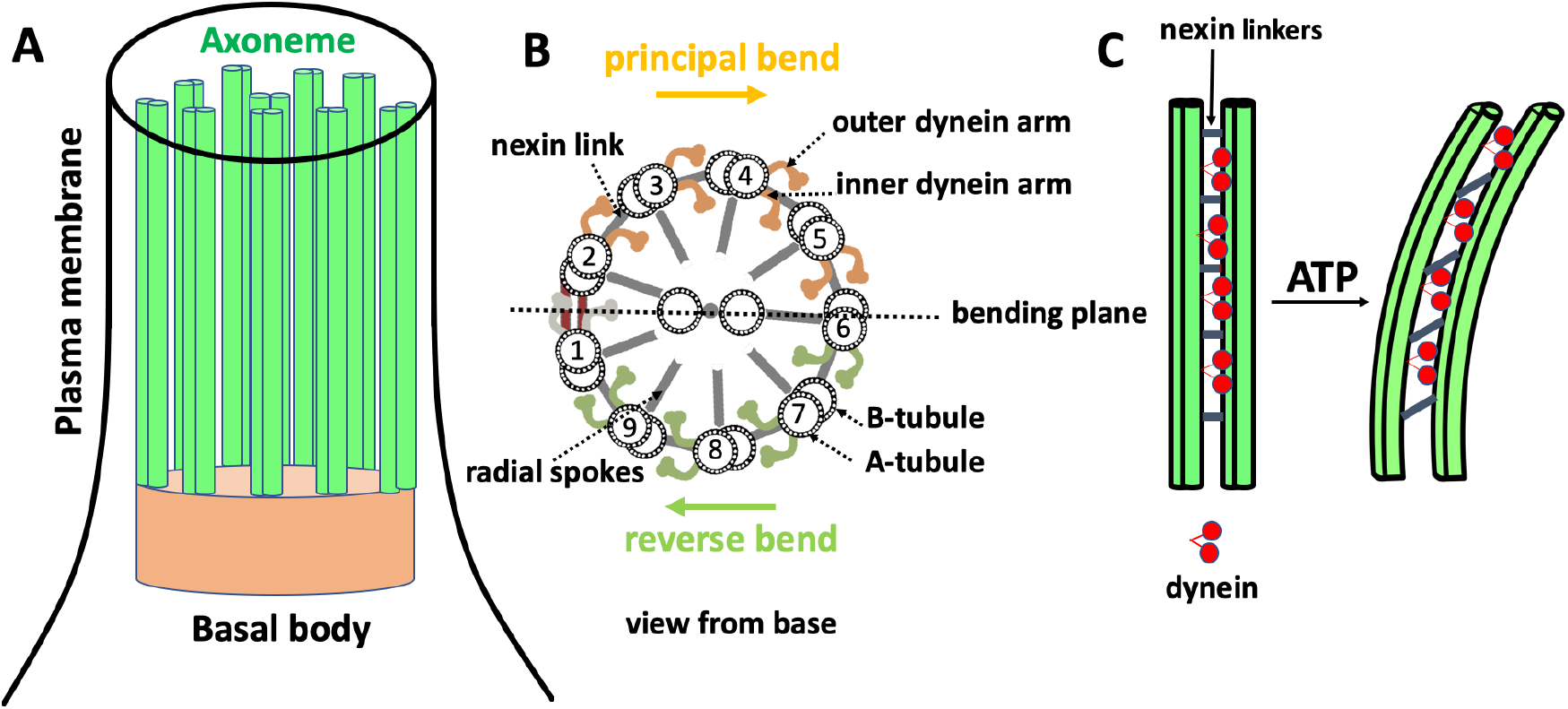
Structure of cilia/flagella. A) Cilia/flagella have a cylindrical architecture composed of axoneme, plasma membrane and basal body. B) Axoneme has a microtubule-based structure formed of nine microtubule doublets at the periphery and two microtubule singlets at the center which are associated with protein complexes such as radial spokes, inner dynein arm, outer dynein arm and nexin linkers. There is a mechanical feedback from the bending on the regulation of dynein activity which switches the activity of dyneins on the opposite side of the central pair microtubules^15^. C) In the presence of nexin proteins which cross-link the neighboring doublets, microtubules are not free to slide and active forces generated by dynein molecular motors bend the cilium/flagellum. Nexin linkers play an important role to convert action of dynein motors in microtubule bending^16,17^ and radial spokes ensure that dyneins work together in a highly coordinated manner to generate a regular wave pattern^18^.

Furthermore, microtubules (MTs) are one of the cytoskeletal filaments which, together with actin and intermediate filaments, provide shape and motility in eukaryotic cells and play a crucial role in formation of various cellular structures such as the dynamic asters found in mitotic and meiotic spindles^28–31^. These spindles have a microtubule-based structure, which generate forces by polymerization-depolymerization or transduce forces generated by ATP-driven molecular motors. Encapsulation of cytoskletal filaments and purified motors^32^ is a powerful tool to study the influence of geometrical confinement on self-organization of motor-driven filamentous network in cell-like compartments^33^. The ultimate challenging goal in the growing field of synthetic biology is to build an artificial cell with a self-sufficient energy conversion system to autonomously power the activity of molecular motors.

Both types of motility requires energy in form of ATP. ATP is also common chemical energy source for many different motility mechanisms found in nature^34^. Therefore, an ATP regeneration module is required for a self-sustained motility. With respect to this, we recently reviewed^1^ different strategies for ATP regeneration in the context of bottom-up synthetic biology and we reported on different types of energy functional modules^35,36^. They are utilizing either chemical^35^ or light energy^36^ to convert ADP to ATP. The latter system appears especially promising for coupling with such bottom-up applications which rely only on ATP as an energy carrier and do not involve NADH (NAD(P)H) reductive power, like in the case of motility. The light energy driven ATP regeneration is not novel; for mimicking natural photophosphorylation, a series of artificial systems have been constructed to capture the energy of light and move protons across the membranes^37^. Simple prototypical systems, which combine light-driven proton pumps with the F_O_F_1_-ATP synthase in liposomes have been demonstrated already in the early 70s, the motivation being to develop *in vitro* models for the mechanistic understanding of F_O_F_1_-ATP synthase. By varying several different types of rhodopsins and F_O_F_1_-ATP synthases as well as of the lipid composition, the productivity of these assemblies was improved as shown by us^36^ and other authors^38–41^. Recently, some examples showing coupling of the light-driven ATP regeneration module with energy intensive tasks like CO_2_ fixation^42^, actin contractions^43^ and protein synthesis^44^ were demonstrated. However, there was no study demonstrating self-sustained system showing directed movement driven by ATP.

Similar as in nature, where different types of motility have evolutionary emerged, artificial cells can also implement different motility mechanisms depending on their intended application. Therefore, we tested a possibility to integrate the energy module with two different motility modules. First, we integrated the light-switchable bottom-up assembled ATP regeneration module with a motility functional unit which was isolated from green algae *Chlamydomonas reinhardtii*. The combination of these two modules leads to an artificial system where energy of light is converted into mechanical work in a self-sustained manner. Next, we co-encapsulated the light-to-ATP energy module with microtubules and kinesin-1 molecular motors to generate active stresses in confined cell-like compartments. We observed controlled motor-driven contraction of filamentous network upon light stimulation, indicating the efficiency of the artificial photosynthetic module in providing self-sufficient energy. This integrated system is potentially applicable in ATP-dependent motility assays, which aim to reconstitute motor-driven motion and force generation inside a synthetic cell, with the challenging goal of achieving light-controllable switch between motile and immotile state of an artificial cell.

## Results

### Light-driven energy module

We engineered light-switchable photosynthetic liposomes (~150 nm in diameter) as energy modules to generate ATP under illumination. To convert light into ATP, we co-reconstituted two purified transmembrane proteins namely, Bacteriorhodopsin (bR) and EF_O_F_1_-ATP synthase from E. coli. Upon illumination, Bacteriorhodopsin pumps proton into the vesicle’s interior, establishing a proton motive force that drives ATP synthase to catalyze the conversion of ADP to ATP (Figure 2A). Both enzymes are isolated according to the procedures earlier described by Ishmukhametov et al.^45^ and Oesterhelt^46^. Their purity is checked by SDS-PAGE analysis (SI, Figure S1). bR is reconstituted in the form of membrane patches to avoid material loss during solubilization. We recently showed that^36^ the usage of monomeric, detergent-solubilized bR for reconstitution is also possible and will lead to a functional ATP regeneration module. However, an almost uniform orientation of bR in phosphatidylcholine (PC) liposomes could only be achieved when using bR in the form of membrane patches (SI, Figure S2). Both enzymes are co-reconstituted into preformed PC unilamellar vesicles using Triton X-100 as detergent, similar to the method described by Fischer and Gräber^47^ for ATP synthase reconstitution. Liposomes are prepared using the extrusion method. The size of vesicles before and after reconstitution is checked using Dynamic Light Scattering (DLS). DLS data confirm an average vesicle diameter of ·~150 nm (SI, Figure S3). Separate assays for each transmembrane protein further prove their functionality. bR proton pumping and intravesicular acidification is detected using pyranine as an internal probe (SI, Figure S4). The activity of ATP synthase is determined in an acid-base transition experiment (SI, Figure S5). ATP production in the co-reconstituted module under illumination is measured using the luciferin/luciferase assay (Figure 2B). The achieved maximal rate of 4.5 μmol ATP (mgEF_O_F_1_)^-1^min^-1^ is high compared to the literature work. An overview of different light-driven and chemically-driven ATP regeneration modules and their ATP production rates can be found in our recent article^1^. We mainly attribute this comparably high efficiency of ATP production to the increased amount of bR reconstituted in our system, aiming for a theoretical ratio of 1 ATP synthase and 96 bR molecules per liposome with an almost uniform direction. In most of the literature work^1,37,39,41,48^ significantly lower bR concentrations were used for co-reconstitution. Only Racker and Stoeckenius^49^ as well as Choi and Montemagno^50^ applied bR concentrations of similar magnitude. Moreover, we found out that the vesicle preparation method has significant impact on the performance of the ATP module. Using dialysis liposomes instead of vesicles produced by extrusion, led to roughly a 50% decrease in activity. Furthermore, Dithiothreitol (DTT) has a positive effect on the ATP production rates. Only 43% of activity remained in the absence of DTT (SI, Figure S6). DTT is known to prevent oxidation and thus to preserve proteins in their functional form. In addition, DTT can contribute to changes in membrane potential, as shown in Ref.^51^. Finally, all experiments have been done with ultra pure ADP (>99,9%). Commonly offered ADP salt is highly contaminated with ATP (>2%). Due to the high ADP concentration in the present measurements (1.6 mM), this would lead to a high ATP concentration at the beginning of the experiments, which could compromise the activity of ATP energy module.

**Figure 2:**
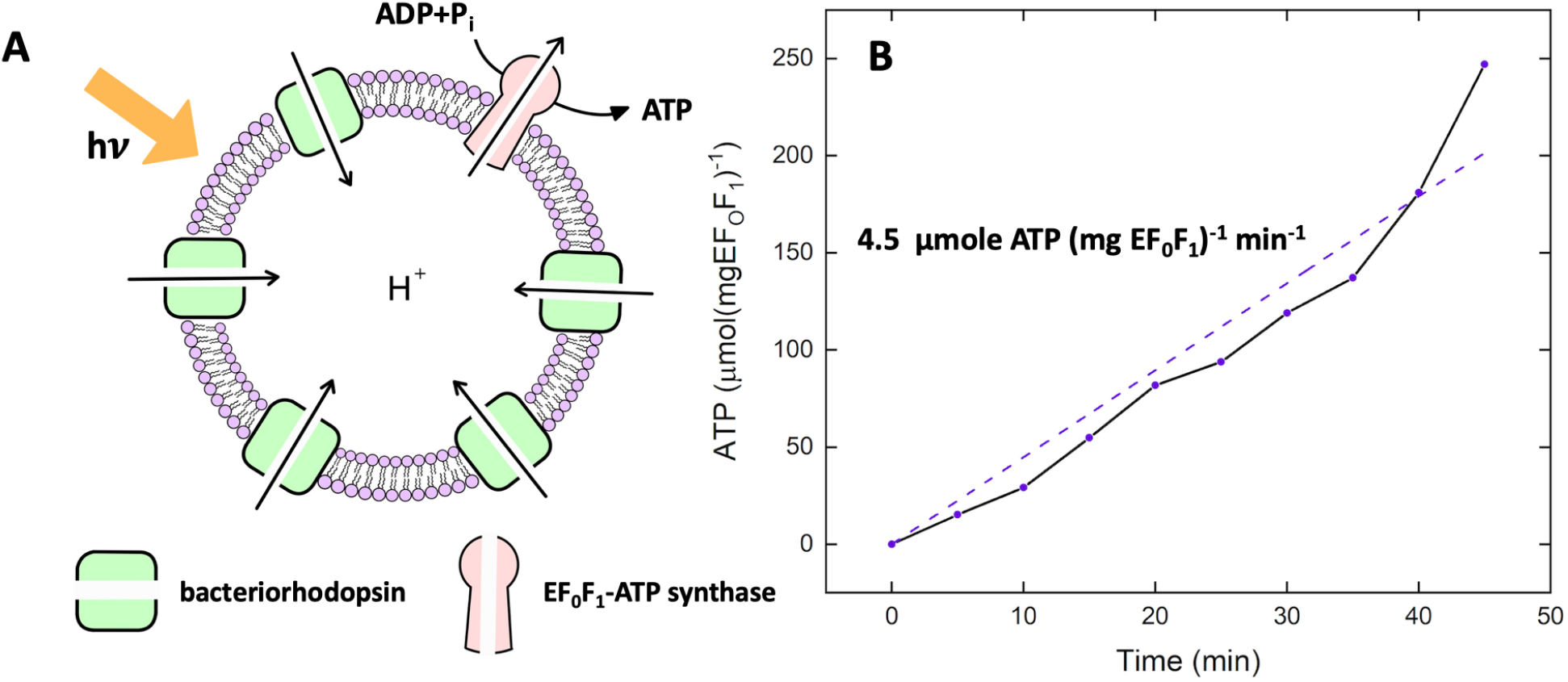
Light-driven ATP production. A) Schematic representation of light-driven ATP synthesis in lipid vesicles. EF_O_F_1_-ATP synthase uses the electrochemical gradient generated by bacteriorhodopsin to synthesize ATP from ADP and P_i_. B) Measurement of light-driven ATP production over time. The maximal rate of 4.5μmol ATP (mgEF_O_F_1_)^-1^min^-1^ is determined by linear regression. Experiments are performed at room temperature with the HMDEKP buffer as the inner solution (30 mM HEPES-KOH, 5 mM MgSO_4_, 1 mM DTT, 1 mM EGTA, 50 mM potassium acetate, 1% (w/v) PEG, pH=7.4). The same outer solution was adjusted with 5 mM NaH_2_PO_4_, 2 mM MgCl_2_, 1 mM DTT and 810μM ADP. [P_i_]= 5 mM, [lipid]=0.022 mg/mL, [EF_O_F_1_] = 2.6 nM, [bR]= 160 nM and ΔΨ= 143 mV is the membrane potential (the outer potential minus the inner potential). Proteins were reconstituted with 0.8 % Triton.

### Integration of energy module with motility module

#### Reactivation of axonemes with pure ATP

In this part, we aim to integrate the light-driven energy module with the motility module, namely the flagella isolated from green algae *C. reinhardtii*^52^ (see Figure 3A-B and Materials and Methods). In the first step, we characterized the activity of the isolated flagella (~10 μm in length) using commercially available pure ATP^53^. Upon mixing ATP with demembranated flagella (axonemes), ATP powers dynein molecular motors that convert chemical energy into mechanical work by sliding adjacent microtubule doublets relative to each other (Figure 1C). However, due to mechanical constraints, MT doublets can not slide freely. Instead, sliding is converted into rhythmic bending deformations that propagate along the contour length of axonemes at a frequency that depends on ATP concentration (Figure 3C-E and SI Videos 1-2). We did not observe beating activity for ATP concentrations below the critical value of [ATP]_critical_ =60 μM^53,54^, suggesting that a minimum number of active molecular motors are required to generate rhythmic motion in axonemes. The critical beat frequency was *f*_critical_ ~14.6 Hz. Above [ATP]_critical_=60μM, the beat frequency increases with [ATP] following a modified Michaelis-Menten kinetics, which predicts a plateau with a linear onset at small values of [ATP]: *f* = *f*_critical_ + *f*_max_([ATP] - [ATP]_critical_)/(*K_m_* +([ATP] - [ATP]_critical_)), with *f*_max_ = 73.75 Hz and *K_m_*=295.8 μM^53,55^.

**Figure 3:**
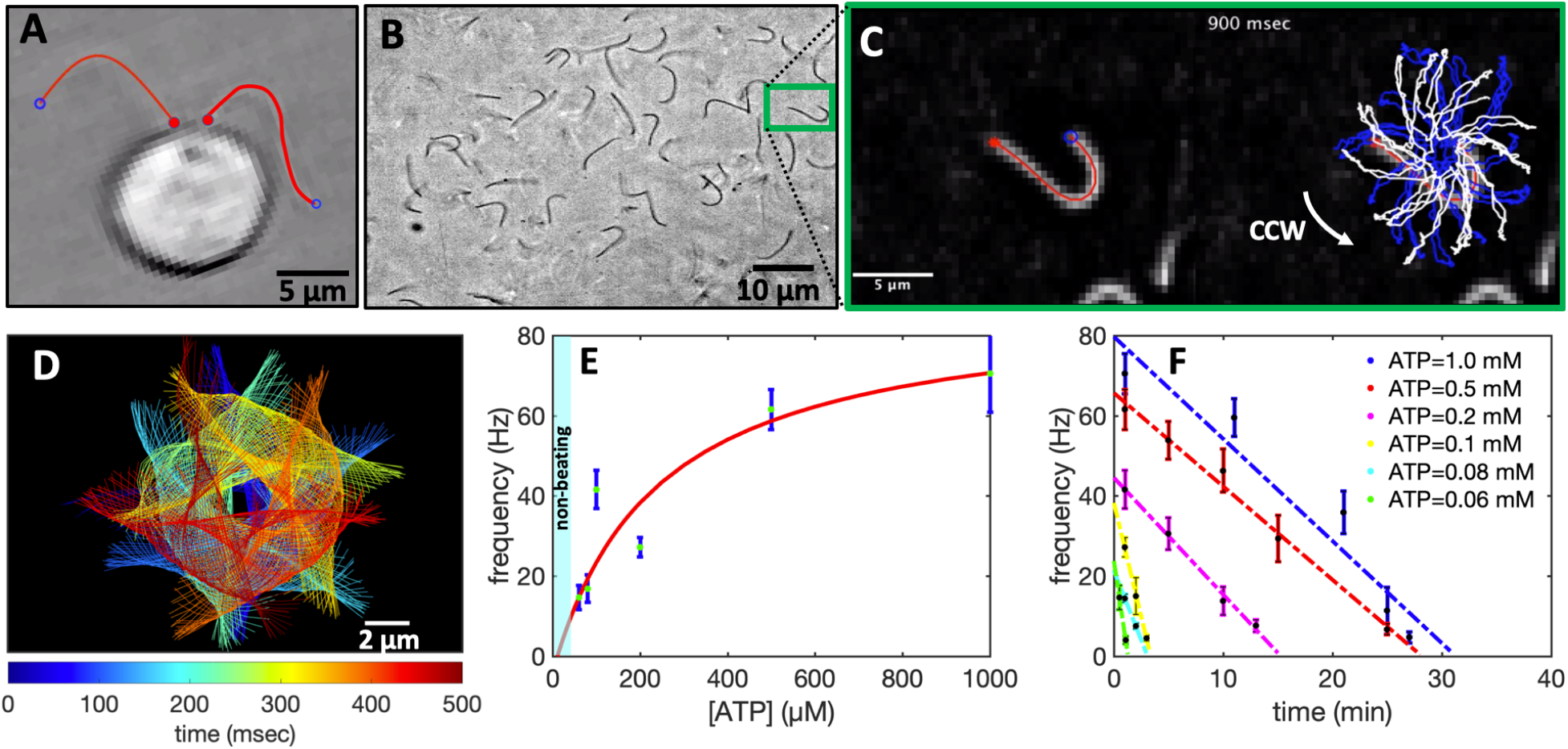
Experiments with pure commercial ATP. A) Snapshot of a *C. reinhardtii* cell with its two flagella. B) Isolated and demembranted flagella are reactivated with pure ATP. C) Swimming trajectory of an exemplary axoneme beating at 18 Hz with [ATP]=80 μM. D) Color-coded time projection of the axoneme in panel C shows the circular swimming path. E) Mean beat frequency as a function of ATP concentration. Solid red line is a least square fit to the modified Michaelis-Menten relation (see text). The critical minimum ATP concentration required to observe axonemal beating was [ATP]_critical_=60 *μ*M. F) Beat frequency decays over time at a rate that depends on ATP concentration. In the presence of 1 mM ATP (blue line) in a 10 *μ*l channel, axonemes beat for 32 minutes at a decreasing beating frequency that allows us to estimate the averaged ATP consumption rate of 0.31 nmole/min. This rate depends on ATP concentration and decreases to 0.25 nmole/min for [ATP]=0.1 mM (yellow line). Error bars are mean ± standard deviation (*N* = 7).

Axonemes consume ATP and beat at a frequency that decays over time until they stop beating (Figure 3F). The rate of ATP consumption depends on both ATP and axoneme concentration. In our experiments, we estimate to have 6 × 10^5^ axonemes in 10 μL of reactivation solution. At an ATP concentration of 1 mM, axonemes stop beating after 32 min, resulting in an estimated ATP consumption rate of 0.31 nmole/min. Assuming that all 6 × 10^5^ axonemes in the chamber are active and consume ATP, we calculate that 5 × 10^6^ ATP molecules/sec are consumed by a single axoneme. Given the mean beat frequency of 50 Hz, we estimate that ~ 10^5^ ATP molecules are consumed in one beating cycle^56^. This averaged consumption rate estimated from our experimental data is comparable to the bulk measurements in the sea urchin sperm^56^ but lower than the value of 2.3 × 10^5^ ATP/beat measured at the single-axoneme level by Chen et al.^55^. This discrepancy can probably be related to the assumption that 100% of axonemes are reactivated, which is not the case even under ideal experimental conditions. Normally, the isolation and demembranation process results in a mixed population of active, non-active, and fragmented axonemes and a 100% reactivation is never achieved. Therefore, single-flagellar experiments, such as those performed by Chen et al.^55^, are more accurate in determining ATP consumption rates.

#### Reactivation of axonemes with light

Next, we combined the light-driven ATP generation module with isolated and demembranted axonemes, as schematically shown in Figure 4A. We first illuminated the energy module for various time intervals between 0 to 45 min before mixing the functionalized vesicles with axonemes. Higher ATP concentrations are produced by illuminating the energy module for longer periods. While illumination with a 5W microscope light generates up to 213 μM ATP, a 50W white light LED lamp produces up to 330 μM ATP after 45 min of illumination (Figure 4B-C). Synthesized ATP reactivates axonemes (SI, Videos 3 and 4) at a frequency that depends on [ATP] (SI, Video 5). The mean beat frequency as a function of [ATP] follows the Michaelis-Menten scaling with *f*_max_ = 50.80 Hz and *K_m_* = 7.14 μM (Figure 4D). Interestingly, we observed that illumination with the microscope light for 1 min, corresponding to 1 μM ATP, was sufficient to reactivate axonemes at a beat frequency of around 22 Hz. This is rather surprising because the minimum critical ATP concentration required to reactivate axonemes in our pure ATP experiments was 60 μM (Figure 3E). We attribute this discrepancy to several factors: 1) In the vicinity of axonemes, ATP is constantly synthesized and subsequently consumed by dynein molecular motors. ATP is known to inhibit the activity of ATP synthase and local consumption of ATP by axonemes can enhance the rate of ATP production. Therefore, the ATP concentration in the presence of axonemes might be slightly higher. 2) Functionalized vesicles may accumulate/adhere to the demembranted axonemes resulting in higher ATP concentrations around them. Our experiments with fluorescently labeled vesicles did not confirm any significant accumulation of vesicles along the entire contour length of axonemes, but we occasionally observed attachment of vesicles to a part of axonemes (SI, Video 6). 3) The last but most important factor is ADP. In contrast to the experiments with pure ATP, experimental system with axonemes and energy module contains a significant amount of ADP (1.6 mM). According to the literature^57–59^, ADP can bind to non-catalytic sites of dynein motors, enhancing the overall energy efficiency of chemical to mechanical energy transformation (Figure 5A-B). In fact, the activating role of ADP in the regulation of on-off switching of dynein arms in flagellar motility has been the subject of several studies in the past years^60–62^. To confirm the activating effect of ADP, we repeated our pure ATP experiments with 1.6 mM ADP. Remarkably, we observed reactivation of axonemes even at a very low ATP concentration of 0.1 *μ*M (Figure 5C). Furthermore, in the presence of ADP, ATP consumption rate was much lower. This is shown in Figure 5D for the fixed ATP concentration of 60 μM with 1.6 mM ADP, where in comparison to the experiment without ADP, axonemes beat at higher frequencies and for a longer period of time.

**Figure 4:**
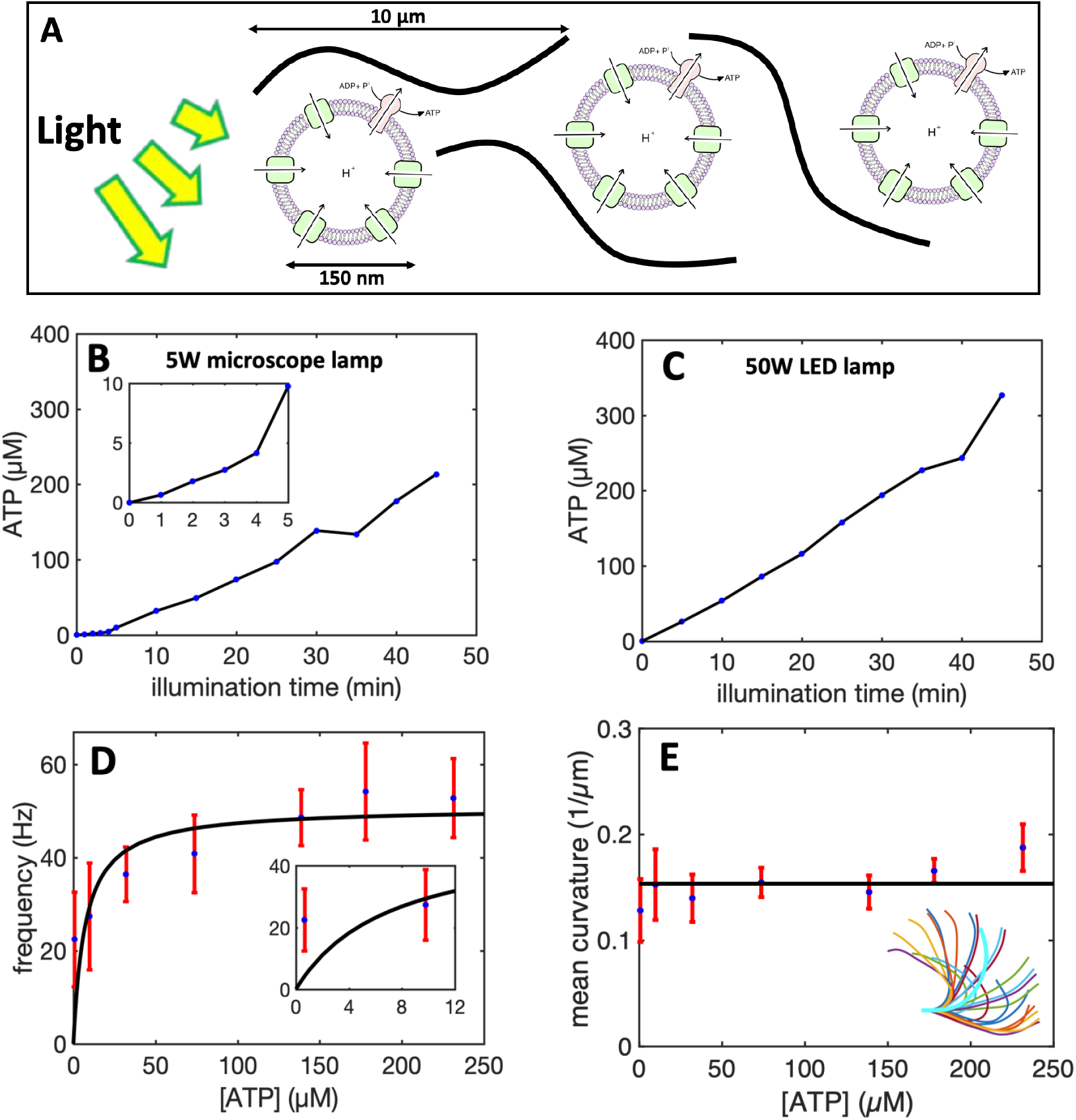
Integration of the motility module with the light-switchable energy module. A) Schematic presentation of isolated flagella mixed with energy module. B) Functionalized liposomes are illuminated for different times, generating ATP concentrations up to 213 μM under illumination with a 5W microscope lamp. C) Higher ATP concentrations up to 330 μM was produced under illumination with a 50W LED lamp. Both light sources are located 25 cm away from the sample. Inset shows ATP production in the time interval 0 to 5 min of illumination. D) Axonemes beat faster at higher ATP concentrations produced by longer illumination of energy module under microscope light. Inset shows that axonemes beat even at small ATP concentrations below 10 μM. E) Static curvature of the axonemes, defined as the curvature of the mean shape averaged over one beating cycle (arc-shaped filament with cyan color), does not significantly depend on ATP concentration. The black line shows a linear fit with the offset of ~0.16 *μ*m^-1^ and slope of zero. For each data point in panels D and E, frequencies of 10 axonemes are measured to calculate the mean and standard deviation.

**Figure 5:**
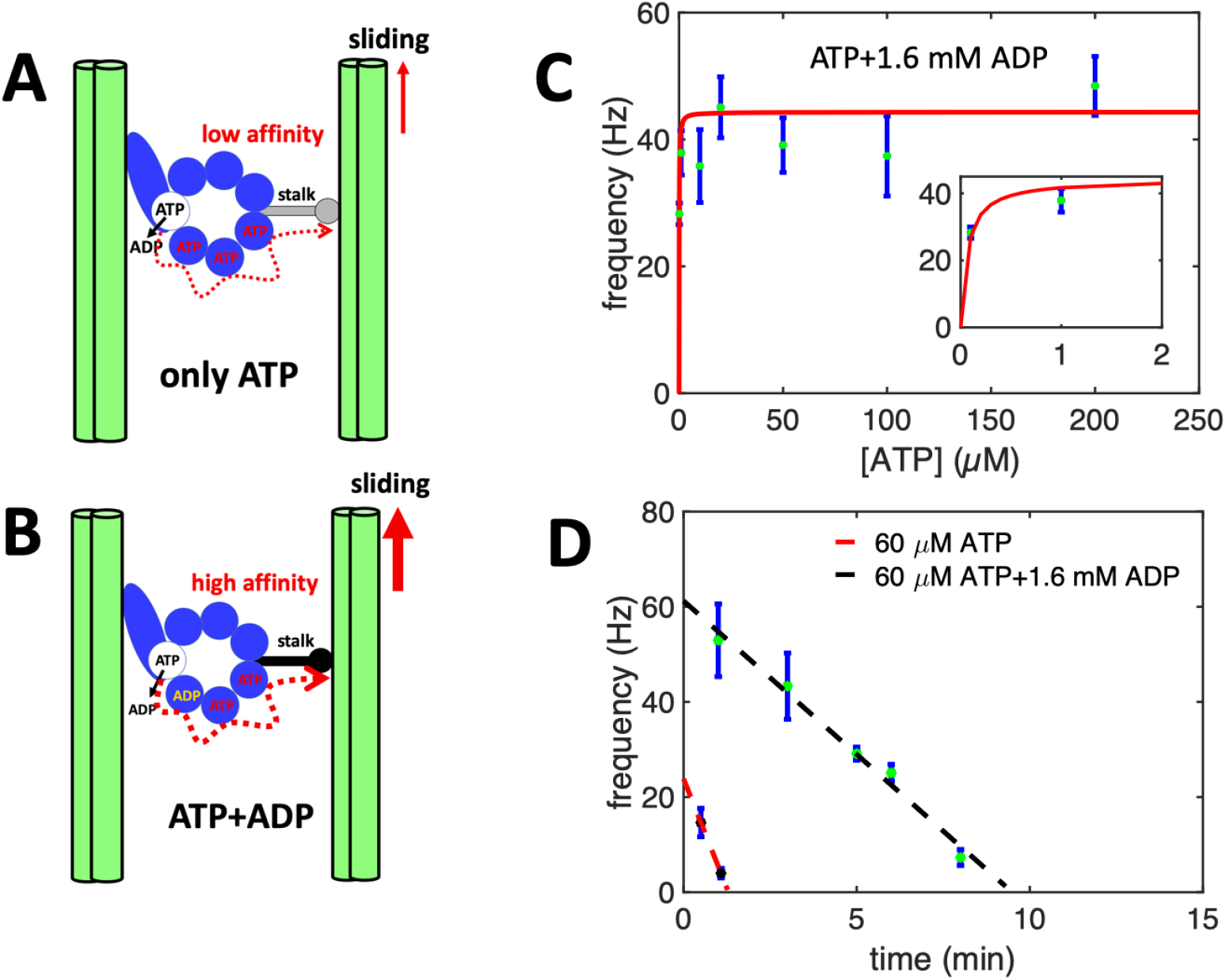
The activating role of ADP. A-B) A hypothetical mechanism introduced in Refs. 57,63 illustrating the regulatory effect of ADP on the binding affinity of dynein to the B-subtubule of the outer MT doublets. Hence, in the presence of ADP which binds to a non-catalytic site, dynein is more efficient in generating the sliding force. C) Pure ATP experiments supplemented with 1.6 mM ADP confirm the activating role of ADP at low ATP concentrations. Note that axonemes are reactivated even at a very low ATP concentration of 0.1 *μ*M. D) Comparison of two sets of experiments with and without ADP at fixed ATP concentration of 60 *μ*M. While without ADP, axonemes stop beating after 2 min, with 1.6 mM ADP, axonemes beat with higher frequencies and are active for a longer time.

ATP consumed by axonemes is replenished by the continuous microscope illumination while we image the sample, establishing an energy production-consumption cycle. Once the microscope light is turned off, the time required for the synthesized ATP to be hydrolyzed depends on both ATP and axoneme concentrations. Exemplary, in a 10 μL solution with 6× 10^5^ axonemes, 60 *μ*M ATP which is produced in a 10 min pre-illuminated energy module, will be consumed in ~10 min. This is consistent with Figure 5D with 60 *μ*M commercial ATP and 1.6 mM ADP. Once ATP is depleted, turning on the microscope light for 1 min generates ~1 *μ*M ATP, which is sufficient to reactivate axonemes but at a lower frequency of ~20 Hz.

Figure 6A-C shows exemplary oscillatory motion of an axoneme in response to ATP generated by the light-driven energy module which is pre-illuminated with the microscope light for 45 minutes (SI, Video 4). We observed bending deformations propagating at a frequency of up to 72 Hz from the basal end toward the distal tip (Figure 6D-E). To analyze the oscillatory motion of axonemes, we first tracked the filaments using the gradient vector flow technique^64,65^ (Figure S7 and Materials and Methods) and quantified the curvature waves using the Frenet equations in a plane^66^:

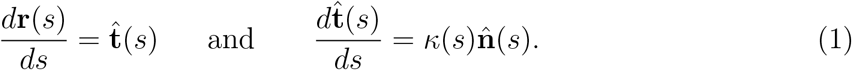

**Figure 6:**
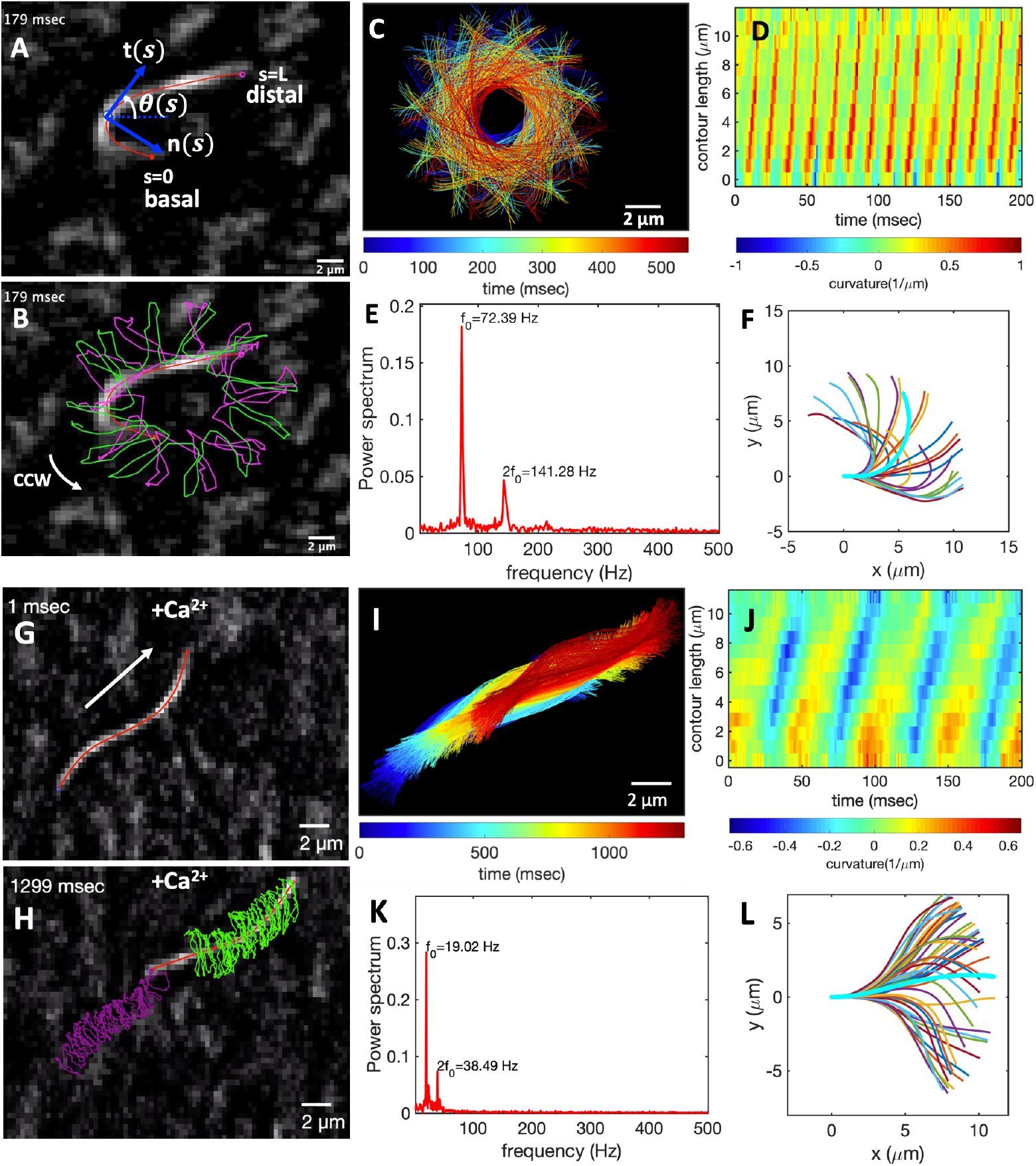
Light-driven reactivation of axonemes with and without calcium. (A) An actively beating axoneme fueled by 213 μM ATP produced during pre-illumination of the energy module for 45 min by microscope light (SI, Video 4). (B) As axoneme beats with frequency of 72 Hz, it rotates CCW with slower frequency of 2 Hz. Magenta and green trajectories show the traces of distal and basal ends of the axoneme, respectively. (C) Color-coded time projections of the beating axoneme showing the circular swimming path. (D) Curvature waves propagate along the contour length from the basal toward the distal end. (E) Power spectrum of curvature waves shows dominant peaks at *f*_0_ = 72 Hz and at second harmonic 2*f*_0_. (F) Configurations of the axoneme at different time points are translated and rotated such that the basal end is at (0, 0) and the orientation of the tangle vector at the basal end is in the x direction. Static curvature of this axoneme is ~0.2 μm^-1^ (G-L) A separate experiment with 1 mM CaCl_2_ which reduces the static curvature of the axoneme to 0.01 μm^-1^ (compare filaments with cyan color in panels F and L). Thus, the axoneme swims in a straight trajectory (compare panels C and I) utilizing ~1 μM ATP synthesized by energy of microscope light without 45 min pre-illumination step. At such a low ATP concentration, axoneme beats at a slower frequency of 19 Hz (SI, Video 7).

Here 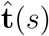 is the unit tangent vector to the axoneme, 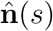 is the unit normal vector, and *κ*(*s*) is the curvature (see Figure 6A). We define *θ*(*s*) as the angle between the tangent vector at contour length *s* and the *x*-axis, then *κ*(*s*) = *dθ*(*s*)/*ds*, which is plotted in Figure 6D. Furthermore, considering reactivated axonemes as an example of a self-sustained biological oscillator, it is possible to perform a principal mode analysis^66^ and define a phase which facilitates a quantitative dynamic analysis of axonemal shape. The interested reader is referred to Materials and Methods and Figure S8.

Finally, to characterize the mean shape of axonemes, which is a circular arc (the filament in cyan color in Figure 6F), we translated and rotated configurations of an actively beating axoneme such that the basal end defined as s = 0 is at position (0, 0) and the tangent vector at the basal end is orientated in the 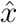 direction (Figure 6F). Figure 4E illustrates that curvature of the mean shape of axonemes does not depend significantly on ATP concentration^53^. This static curvature (~0.16 μm^-1^) which leads to an asymmetric waveform and causes a circular swimming trajectory of axonemes (Figure 6B-C), is comparable to the values obtained in our pure ATP experiments with and without ADP (see Figure S9 and Ref.^53^) and can be reduced by adding calcium ions to the reactivation buffer^67^. Calcium ions are known to play a crucial role in shaping and controlling flagellar waveform^67,68^. A calcium-responsive protein at the interface between radial spoke RS1 and inner dynein arm IDAa is calmodulin (see Figure 1B and Figure 5 in Ref.^69^). Electron cryotomography (cryo-EM) data^69^ show that structure of calmodulin undergoes a Ca^2+^-dependent conformational change which could affect the RS1-IDAa interaction, therefore modulating the flagellar beat from an asymmetric to a symmetric waveform. Figure 6G-L shows an exemplary axoneme with a symmetric waveform in which the static curvature is reduced to 0.01 μm^-1^ by addition of 1 mM CaCl_2_. This axoneme with nearly ten times reduced static curvature swims in a straight line (SI, Video 7). We emphasize that beat frequency, static curvature and amplitude of curvature waves are three important factors which determine the swimming trajectory of an axoneme in the ambient fluid and variations in these parameters directly affect the swimming dynamics (see Materials and Methods, Figure S10 and Video 8 for simulations of swimming trajectory).

### Light-driven contraction of MTs/kinesin-1 network

We further tested the suitability of light-switchable energy module for encapsulation of *in vitro* motility assays constituted of microtubules and force-generating molecular motors. This assembly under illumination provides an energetically autonomous system that has the potential to advance the development of synthetic cells (Figure 7A).

**Figure 7:**
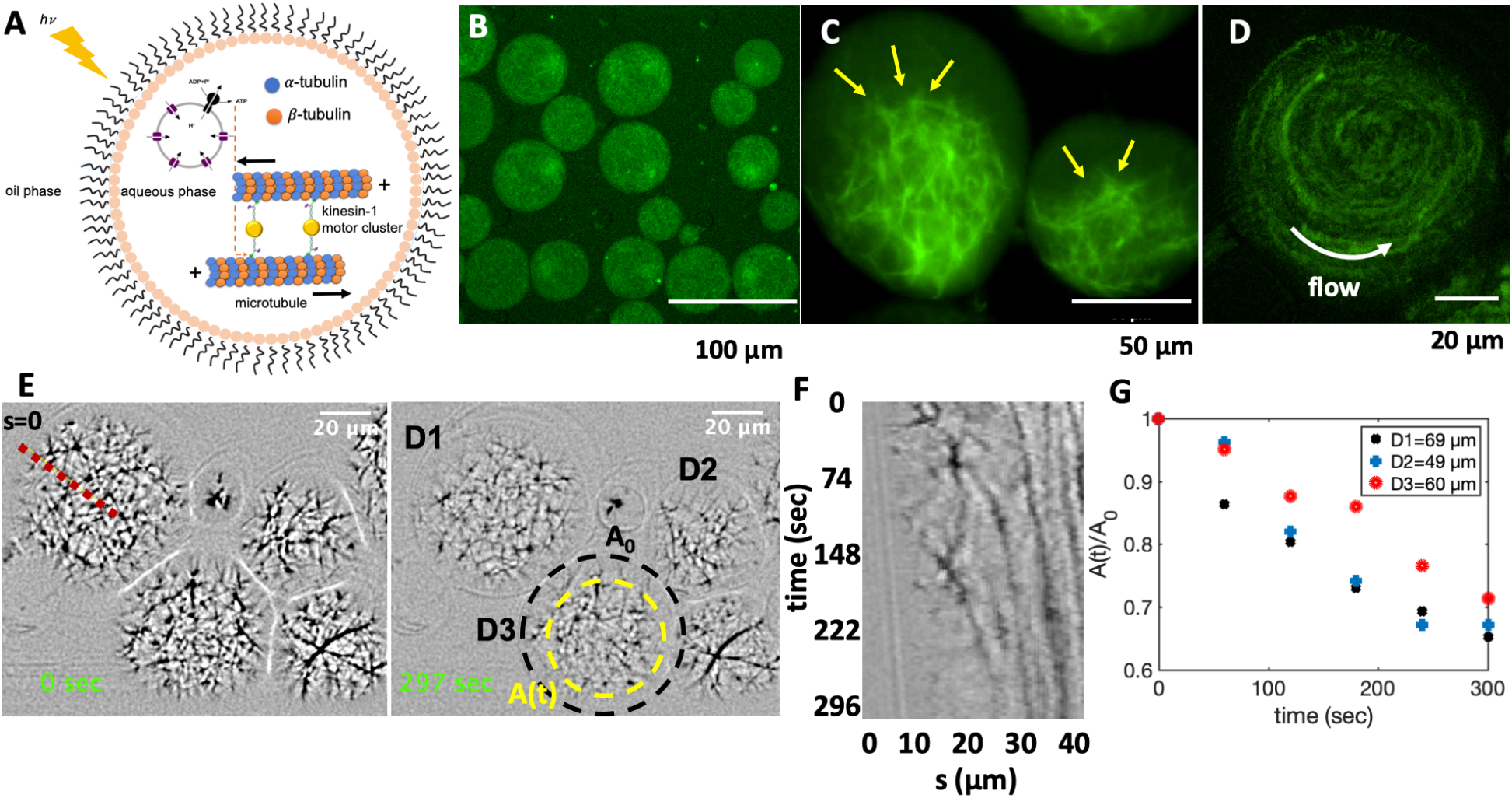
Cell-like confinement of MTs, kinesin-1 molecular motors and light-switchable energy module. Kinesin-1 is a dimer with two heads which forms cluster through streptavidin. A) Schematic representation of the MTs/kinesin1 network co-encapsulated with functionalized vesicles inside water-in-oil droplets. Upon illumination, synthesized ATP provides fuel for the kinesin-1 molecular motors, which are plus-end directed motors that exert contractile stresses by sliding MTs relative to each other. B) MTs/kinesin-1 network shows a relatively uniform distribution shortly after encapsulation. C) Snapshots of network contraction after 40 mins. The yellow arrows indicate network contraction. D) Rotational flows observed in some of the droplets during network contracts. E) Snapshots of contractile active network at two different time points. The yellow circle shows the area covered by the network after ~5 min. F) Space-time plot showing network contraction along the red dashed line in panel E. G) Relative reduction of network area over time for three droplets shown in panel E. A_0_ is the initial area of the network before contraction, as marked with a black circle in panel D.

We verified the activity of the motor-driven filamentous network by characterizing its activity in water-in-oil droplets of various diameters. Figure 7 and supplemental Video 9 show the contraction of MTs/kinesin-1 network mixed with the pre-illuminated photosynthetic energy module and encapsulated in a microfluidic device (see Materials and Methods and Figure S11 for the design of the set up). We observed a relatively uniform distribution of the network immediately after encapsulation, as shown in Figure 7B. Over time, ATP produced in pre-illuminated energy module drives the activity of molecular motors allowing them to cross-link and slide neighboring microtubules against each other. This results in a net force that contracts the network^70^. A snapshot of the contracted network after 40 mins is shown in Figure 7C. Microtubule-microtubule sliding also causes rotational flows within droplets, which were occasionally observed in our experiments, as shown in Figure 7D and discussed in Ref.^71^ (SI, Video 9). We quantified the contractile activity of the system by analyzing the relative reduction in the network area over time. Figure 7E-G shows that network contraction within droplets of different diameters occurs at similar time-scales. We note that discontinuous illumination of the sample by microscope light every 5 seconds during imaging, produces ATP which compensates consumption of ATP by molecular motors. We verified this circulating energy-production and consumption in the system by a separate experiment where we confined the MTs/kinesin-1 network in a milli-fluidic device with rectangular cross-section (30 mm×1.5 mm×0.1 mm). The aim is to compare the contractility of the network in areas illuminated by discontinuous microscope light, where ATP is consumed and produced, versus non-illuminated areas, where ATP is only consumed but not replenished by light. Figure 8 shows that initially the active network fills the entire 3D volume of the channel. Kinesin-1 motors consume ATP synthesized in pre-illuminated energy module, generating active stresses which results in network contraction (Figure 8B-D). As kinesin-1 motors consume ATP, discontinuous microscope illumination compensates ATP consumption and motor-driven network contraction reaches its maximum value of 38.6% within 60 min, as displayed in Figure 8B-D and supplemental Video 10. ATP production is maintained only in areas illuminated by the microscope light, causing stronger network contraction compared to non-illuminated regions in the channel. We observed a reduced contractility of about 10% in the adjacent volume near the region illuminated by the microscope light, as shown in Figure 8E. These experiments suggest that the application of patterned illumination in a filamentous network can impose different contraction-maps with spatial gradient. It also demonstrates the efficiency of the light-driven energy module for controlled and localized ATP production, and highlights the potential application of this system in the field of synthetic biology with the ultimate goal of building an artificial cell.

**Figure 8:**
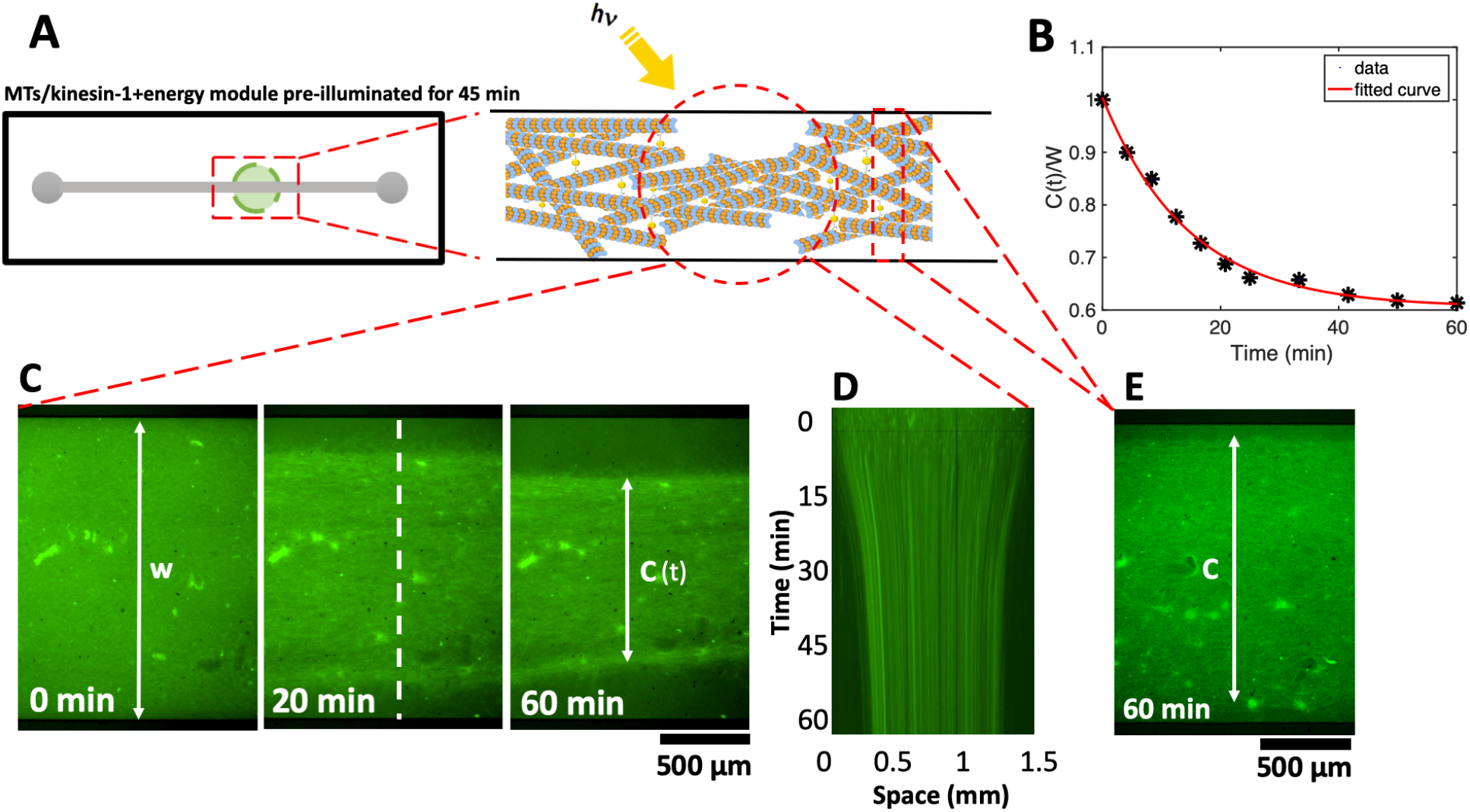
Photo-stimulated contraction of MTs/kinesin-1 network inside a milli-fluidic device (SI, Video 10). A) Schematic representation of the milli-fluidic device highlighting the area illuminated with discontinuous microscope light. Taxol-stabilized MTs are mixed with kinesin-1 motors and a pre-illuminated energy module prior to injection into a milli-fluidic channel. B-C) Over time, as available ATP in the energy module is consumed by forcegenerating kinesin-1 motors, discontinuous microscope illumination compensates for ATP consumption and filamentous network contracts up to 38%. Quantitative analysis of the width of the contracted network in the illuminated region shows an exponential decay over time. The initial width of the network is W= 1.5 mm. D) Space-time plot demonstrating the network contraction along the white dashed line drawn in part C. E) In the non-illuminated areas, where ATP is only consumed but not replenished, we monitored a reduced contractility of up to 10% after 1 hour.

## Discussion

Our experiments demonstrate that light-switchable photosynthetic liposomes can drive ATP-dependent activity of axonemal dyneins, which serve as tiny protein machinery to convert chemical energy into mechanical work in the form of a rhythmic beating pattern. We used light energy to dynamically synthesize ATP, allowing us to control beating frequency of axonemes as a function of illumination time. We illuminated the functionalized vesicles at two different light intensities: a 5W microscope lamp and a 50W white LED lamp, which can generate up to 213 μM and 330 μM ATP, respectively, after 45 mins of illumination (compare Figure 4B and C). We note that bacteriorhodopsin (bR) has a purple color and therefore absorbs green light (500-650 nm) most efficiently. Since bR has a broad excitation spectrum, proton pumping is also possible using white light or red light for excitation. Our results indicate a nonlinear dependence of ATP synthesis rate on light intensity. To generate ATP, ADP is phosphorylated via the F_0_F_1_-ATP synthase reaction:

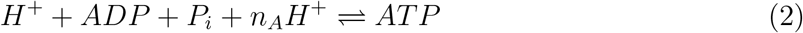

where *n_A_* ~ 2 – 3 is the number of protons transported from the inside to the outside of the vesicle each time the reaction of ATP formation takes place, and *H*^+^ indicates hydrogen ions transferred across the vesicle membrane (Figure 2A). The rate of ATP formation according to Ref.^72^ can be expressed as:

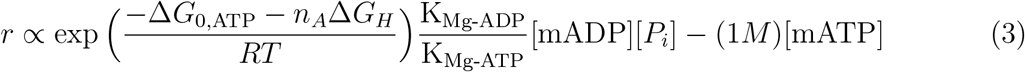

where *R* is the universal gas constant, *T* is the temperature, [mADP], [P_*i*_] and [mATP] are the concentrations of magnesium-bound ADP, P_*i*_ and magnesium-bound ATP and the factor (1*M*) is used to balance the units. K_Mg-ATP_ and K_Mg-ADP_ are the equilibrium dissociation constants for ATP and ADP binding with Mg^2+^ and Δ*G*_0,*ATP*_ = –36.03 kJ/mole is the standard Gibbs free energy for ATP formation. Finally, 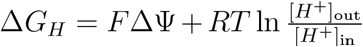, where 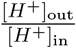 is the ratio of external to internal hydrogen ion concentration, ΔΨ is the membrane potential and F is the Faraday’s constant. The concentration of protons inside the vesicles depends on proton pumping by bacteriorhodopsin, but also on proton pumping by ATP synthase, proton leak across the membrane and buffer capacity inside the vesicles. Only the rate of proton pumping by bacteriorhodopsin can be considered to be approximately linearly dependent on light intensity^73^, but in general the rate of ATP production *r* will be nonlinearly dependent on the light intensity. Some experimental studies on similar systems^48,74^ have confirmed nonlinear dependence of ATP synthesis rate on light intensity and have even shown saturation at higher light intensities, reminiscent of Michaelis-Menten-type kinetics.

In our reactivation experiments with the energy module, we observed that the minimum ATP concentration required to reactivate axonemes is much smaller than the critical value in control experiments with pure commercial ATP (compare Figures 3E and 4D). Multiple factors potentially account for this discrepancy: 1) ATP is known to be one of the inhibitors of ATP synthase activity, and local consumption of ATP by axonemes can enhance the conversion rate of ADP to ATP, resulting in locally higher ATP concentrations near axonemes. To verify the inhibitory effect of ATP, we coupled the light-driven ATP module with a metabolic module by combining the light-driven ATP production with the consumption of glucose, as schematically shown in supplemental Figure S12A. As a proof of concept, a relatively simple metabolic reaction was chosen: Hexokinase converts glucose (G) to glucose-6-phosphate (G-6-P) under the consumption of one molecule of ATP^49^. Our measurements confirm that in the metabolically coupled system, the ATP production rate is 18% higher than the detected ATP production rate in the control experiments; see Figure S12 B-C and Materials and Methods. 2) The attachment of functionalized vesicles to the demembranted axonemes can generate higher ATP concentrations around the axonemes. Although we did not observe vesicle accumulation along the entire contour length of axonemes, we occasionally observed vesicles attaching to some parts of axonemes (SI, Video 6). 3) The most important factor which explains the discrepancy is the presence of ADP. In contrast to the experiments with pure ATP, the experimental system with light-to-ATP energy module was additionally supplemented with ADP at a concentration of 1.6 mM. Remarkably, our experiments with pure ATP supplemented with 1.6 mM ADP confirm the activation role of ADP in reactivation of axonemes at low ATP concentrations below 60 μM (see Figure 5C). We observed beating activity even at 0.1 μM ATP which is much smaller than critical value of 60 μM in pure ATP experiments without ADP. According to the literature, ADP plays a crucial role in axonemal motility1^5,57,60,75^. For example, Inoue and Shingyoji^57^ investigated the influence of ADP and ATP on regulation of dynein activity to produce inter-microtubule sliding in flagella. Dynein motors have four ATP bindings sites in each of their heavy chains (Figure 5A-B). Only one of these sites is catalytic and responsible for the conversion of ATP to ADP while the other three sites are non-catalytic. According to this study^57^, one of these sites can be occupied by ADP. In the absence of ADP, the mean flagellar velocity was lower than with ADP. Moreover, in the presence of ADP, the mean velocity reaches its steady state value faster than in the absence of ADP, suggesting that ADP increases the efficiency of energy transduction. This is consistent with our experiments with ADP in which much lower ATP concentrations were required for axonemal beating. In addition, beating frequency with ATP and ADP reaches saturation at very low ATP concentrations, whereas kinetics with pure ATP is much slower (compare Figure 3E with Figures 4D and 5C). Now, considering that very low ATP concentrations around or below 1 μM ATP (active form is actually ATP-Mg) are required to observe axonemal beating, the question arises whether ADP can bind to more than one of the non-catalytic binding sites in dynein (see Figure 5B).

Self-sustained motility systems that rely on light energy as a primary source of energy (or for *in vivo* applications, chemical energy such as glucose^35^), may have potential applications in the area of synthetic swimmers and targeted drug delivery. In our preliminary experiments, to build a sperm-like swimmer, we attached a cargo (1 μm beads) to the distal end of an axoneme that can be propelled by external illumination (Figure S13 A-B and supplemental Video 11). In light-to-ATP or chemical-to-ATP energy modules, enhanced attachment of functionalized vesicles to the contour length of axonemes could be beneficial to this system^76^ because ATP will be produced locally around axonemes and consumed subsequently by dynein molecular motors (Figure S13 C). These functionalized vesicles attached to an axoneme can also be used as drug carriers^77–79^. Extensive experiments in the future are necessary to investigate the feasibility of these ideas for biomedical applications.

Lastly, we have shown that functionalized artificial liposomes capable of continuous production of ATP in response to light as an external stimulus, can serve as an efficient energy source for *in vitro* microtubule motility assays in which kinesin-1 molecular motors are actively engaged in generating and sustaining active stresses in the network. In these experiments, pre-illumination of the energy module for 45 min generates sufficient ATP needed for motor-driven network contraction. Discontinuous microscope illumination every 5 seconds replenishes the consumed ATP and the MTs/kinesin1 network confined in a milli-fluidic device contracts by up to 38% within 60 min, following an exponential trend (Figure 8B). The encapsulated MTs/kinesin1 network contracts similarly, but in a much faster time scale of ~5 min. Once contracted, the network which is highly cross-linked by kinesin-1 motors and randomly oriented, remains contracted even after the microscope light is turned off and the ATP is depleted. Thus, a light-controllable reversible switch between contracted and relaxed states does not occur in our experiments. We also performed control experiments with commercial ATP, to confirm that a contracted network does not relax back once ATP is completely consumed (see SI, Figure S14 and Video 12 for a 19 hours long experiment with 1 mM pure ATP). Further, the pre-illumination of the energy module is a requirement for the network contraction. Indeed, a MTs/kinesin1 network mixed with a non-preilluminated energy module does not contract, indicating that ATP produced by discontinuous microscope illumination is insufficient to generate critical contractile stresses in the network (SI, Video 13).

Our developed scheme of circulating energy consumption and production could serve as a potential platform to encapsulate constituent elements such as actin, microtubules and various regulatory components inside functionalized lipid vesicles to provide ATP in an optical-controllable self-sustained manner. ATP-driven motor activity in filamentous biopolymer networks is expected to generate active forces that drive morphological deformation in liposomes and further contributes to the challenging goal of bottom-up creation of an artificial cell.

## Material and Methods

### Expression and purification of membrane proteins

Purple membrane was isolated from Halobacterium salinarium (strain S9) as described by Oesterhelt^46^. His-tagged E. coli F_O_F_1_-ATP synthase (EF_O_F_1_) was expressed from the plasmid pBWU13-*β*His in the E. coli strain DK8 (ΔuncBEFHAGDC) and purified by Ni-NTA affinity chromatography as previously described by Ishmukhametov et al.^45^

### Preparation of lipid vesicles for light-driven ATP production

Vesicles were formed by film rehydration method followed by extrusion. 10 mg of dissolved phosphatidylcholine lipid was deposited in a glass vial and solvent was removed using a gentle stream of nitrogen. Thin lipid films were rehydrated in HMDEKP buffer (30 mM HEPES-KOH, 5 mM MgSO_4_, 1 mM DTT, 1 mM EGTA, 50 mM potassium acetate, 1 % PEG, pH 7.4) to a final concentration of 10 mg/mL by vortexing. To transform multilamellar vesicles into unilamellar vesicles, the suspension was subjected to 5 freeze-thaw cycles. Each cycle consisted of freezing in liquid nitrogen, thawing in a 35°C water bath and vortexing for 30 s. Suspensions were extruded 11 times through a 100 nm pore size polycarbonate membrane (Whatman) to form uniform vesicles.

### Co-reconstitution of EF_O_F_1_-ATP synthase and bR

100 μL of preformed vesicles were mixed with 0.1 μM EF_O_F_1_-ATP synthase and 9.9 μM bR in form of membrane patches. 0.8 % Triton X-100 was added under vortexing to partially solubilize the vesicles. After 15 minutes incubation in the dark under gentle shaking, 80 mg of wet SM-2 Bio-Beads were added and the solution was incubated for further 60 minutes under constant shaking in the dark.

### Light-induced ATP production

For measurement of light-induced ATP production, 25 μL of co-reconstituted vesicles were diluted in 250 μL of HMDEKP buffer (30 mM HEPES-KOH, 5 mM MgSO_4_, 1 mM DTT, 1 mM EGTA, 50 mM potassium acetate, pH 7.4) supplemented with 1% (w/v) polyethylene glycol (M_*w*_ = 20 kg mol^-1^) and 1.6 mM ultra pure ADP (Cell Technologies). The reaction was started by illumination with a 50W green LED lamp or a 5W microscope light. Aliquots of 25 μL were taken at the beginning every 1 minutes and later on every 5 minutes from the reaction mixtures and the reaction was stopped by addition of the same volume of trichloroacetic acid (40g/L). The ATP concentration was measured with the luciferin/luciferase assay and calibrated by addition of 10 μL ATP (7.8 μM) after each measurement.

### Isolation and reactivation of flagella with light-driven energy module

Axonemes were isolated from wild-type *Chlamydomonas reinhardtii* cells, strain SAG 11-32b. They were grown axenically in TAP (tris-acetate-phosphate) medium on a 12h/12h day-night cycle. Flagella were isolated using dibucaine,^80,81^ then purified on a 25% sucrose cushion, and demembranated using detergent NP-40 in HMDEK solution supplemented with 0.2 mM Pefabloc. The membrane-free axonemes were resuspended in HMDEK buffer plus 1% (w/v) polyethylene glycol (M_*w*_ = 20kg mol^-1^), 0.2mM Pefabloc, 20% sucrose and stored at −80 °C. To perform reactivation experiments, we thawed axonemes at room temperature and kept them on ice and used them for up to 2 hours. Next, we diluted axonemes in HMDEKP reactivation buffer (HMDEK plus 1 % PEG) which contains light-driven energy module. We infuse them into 100μm deep flow chambers, built from cleaned glass and double-sided tape. Glass slides are treated with casein (2mg/mL in HMDEKP buffer) for 5 min before use to avoid axoneme-substrate attachment. For experiments with calcium, 1 mM CaCl_2_ was added to HMDEKP reactivation buffer.

### Axoneme contour tracking

We imaged reactivated axonemes using phase-contrast microscopy (100X objective, imaging frequency of 1000 fps). To increase the signal to noise ratio, we first inverted phasecontrast images and then subtracted the mean-intensity of the time series. This background-subtraction method increased the signal to noise ratio by a factor of three53. Next, we applied a Gaussian filter to smooth the images. Tracking of axonemes is done using gradient vector flow (GVF) technique64,65. For the first frame, we select a region of interest that should contain only one actively beating axoneme. Then, we initialize the snake by drawing a line polygon along the contour of the axoneme in the first frame (see Figure S7). This polygon is interpolated at *N* equally spaced points and used as a starting parameter for the snake. The GVF is calculated using the GVF regularization coefficient *μ* = 0.1 with 20 iterations. The snake is then deformed according to the GVF where we have adapted the original algorithm by Xu and Prince for open boundary conditions. We obtain positions of *N* points along the contour length *s* of the filament so that *s* = 0 is the basal end and *s* = *L* is the distal side, where *L* is the total contour length of filament. The position of filament at *s_i_* is denoted by *r*(*s_i_*) = (*x*(*s_i_*), *y*(*s_i_*)).

### Simulations with simplified wave form

To investigate the influence of frequency, static curvature and amplitude of the curvature waves on swimming trajectory of axoneme, we performed simulations with a simplified form of curvature waves which is a superposition of a dynamic mode (cosine wave) and a static mode (a circular arc)^52^:

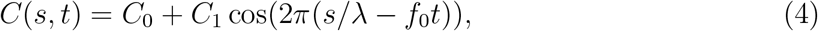

where *C*_0_ is the static curvature, *C*_1_ is the amplitude of the dynamic mode, λ is the wavelength and *f*_0_ is the beating frequency. Minus sign in term —*f*_0_ generates waves that propagate from *s* = 0 (basal end) to *s = L* (distal tip).

As shown in Figure S10A-B, in the absence of a static curvature (*C*_0_ = 0), flagella swims on a straight path and faster beating flagella swims a longer distance. For a non-zero static curvature (*C*_0_ = 0), which is the case for isolated axonemes in our experiments, flagella swims on a circular path, as shown in Figure S10C. Note that in our experiments, adding calcium reduces the static curvature *C*_0_ in a dose dependent manner, approaching to almost zero at 1mM CaCl_2_ concentration (see Figure 6K).

In these simulations, we compute drag force density felt by each flagellum in the framework of resistive-force theory^82^. In this theory, each flagellum is divided to small cylindrical segments moving with velocity **u**_⊥_ and **u**_ǁ_ in the body-frame and propulsive force is proportional to the local centerline velocity components of each segment in parallel and perpendicular directions.

### Mode analysis

We used the Frenet equations to analyze axoneme’s shapes in a plane^66^:

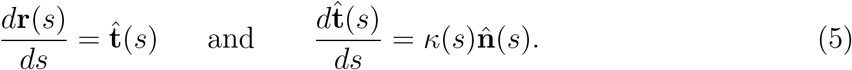

Here 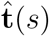 is the unit tangent vector to the axoneme, 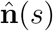 is the unit normal vector and *κ*(*s*) is the curvature (see Figure 6A). We define *θ*(*s*) to be the angle between the tangent vector at contour length *s* and the x-axis, then *κ*(*s*) = *dθ*(*s*)/*ds*. Note that we rotate and translate the axoneme such that point *s* = 0 (basal end) is at position (*x,y*) = (0, 0) and the local tangent vector at *s* = 0 is in the *X* direction, giving *θ*(0) = 0. Following Stephens et al. 66, we performed principal mode analysis by calculating the covariance matrix of angles *θ*(*s*) defined as *C*(*s,s*’) = 〈(*θ*(*s*) — 〈*θ*〉(*θ*(*s*’) — 〈*θ*〉)〉. We then calculated the eigenvalues *λ_n_* and the corresponding eigenvectors *V_n_*(*s*) of this matrix, showing that superposition of 4 eigenvectors corresponding to four largest eigenvalues can describe the axoneme’s shape with high accuracy,

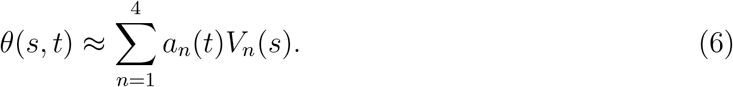

Here the four variables *a*_1_(*t*),..,*a*_4_(*t*) are the amplitudes of motion along different principal components and are given by 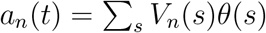.

First, we used data of tracked axonemes obtained by GVF technique^64,65^ to compute *θ*(*s,t*) which is defined as the angle between the local tangent vector and 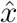 axis (see Figure 6A). Next, we calculated covariance matrix *C*(*s, s*’) of fluctuations in the angle which has only a small number of nonzero eigenvalues. The color map of this matrix presented in Figure S8A shows that *C*(*s, s*’) has an effective reduced dimensionality. As shown by Stephens et al.^66^ for C.elegans, only four eigenvectors *V_n_*(*s*) (*n* = 1,.., 4) corresponding to the first four largest eigenvalues of covariance matrix are enough to describe axoneme’s shape with high accuracy (Figure S8G-I). The first two motion amplitudes *a*_1_(*t*) and *a*_2_(*t*) are shown in Figure S8C, D and the corresponding probability density is presented in Figure S8B (see Materials and Methods). Considering reactivated axonemes as an example of a self-sustained biological oscillator, it is possible to define a phase which facilitates quantitative analysis. To present instantaneous state of an oscillator, we consider the stable limit cycle that forms in *a*_1_ — *a*_2_ plane, and define phase as a monotonically increasing function *ϕ*(*t*) (Figure S8B, F). For a given limit cycle, phase *ϕ* rotates uniformly in the phase space such that *dϕ/dt* = 2*πf*_0_, where *f*_0_ is the autonomous beating frequency of axoneme (Figure S8F).

### Motile bundle solution

The motile bundle solution was obtained by mixing taxol-stabilized microtubules and an active mixture (AM) containing molecular motors kinesin-1 clustered in multimotors configuration by using streptavidin. Kinesin 401 was purified as previously published^83,84^ and the kinesin-streptavidin complexes were prepared by mixing 0.2 mg/mL kinesin 401, 0.9 mM dithiothreitol (DTT), 0.1 mg/mL streptavidin dissolved in M2B (80 mM PIPES, adjusted to pH = 6.9 with KOH, 1 mM EGTA, 2 mM MgCl_2_) and incubated on ice for 15 min.

The AM was obtained by mixing 2.4 mM Trolox, 16.6 μL 3% PEG, 5.5 μL M2B, 3.25 μL DTT (10mM), the oxygen scavenger (0.2 mg/mL glucose oxidase (Sigma G2133), 0.05 mg/mL catalase (Sigma C40)), 0.5 mg/mL glucose, and 4μL kinesin1-streptavidin clusters. The microtubule polymerization mixture (MT) was prepared by mixing 27 μM porcine brain tubulin (1:5 labeled tubulin) in M2B with 5 mM MgCl_2_, 1 mM GTP, 50 % DMSO and 0.3% PEG. This solution is kept in the oven for 30 minutes by 37°C and diluted up to 200 μL with M2B and 7 μM taxol. The final mixture was prepared by mixing polymerized MT, AM and light-driven ATP module in different amounts depending on the experimental set up.

### Encapsulation of MTs/kinesin-1 network assembled with photosynthetic vesicles

The feasibility of the proposed concept, towards building artificial cells, was demonstrated by encapsulation (water-in-oil droplets, w/o) of MTs/kinesin-1 mixed with pre-illuminated ATP module by droplet microfluidic technique (see next section). Sample containing the MTs/kinesin-1 and 45 min pre-illuminated ATP module with volume ratio of 1:20 v/v (ATP module:MTs mixture) was pumped into inner-inlets (aqueous phase 1 and 2), while oil phase (FluoSurf 2% in HFE 7500 or FC40) was pumped into outer-inlet. Micro syringe pumps (CETONI BASE 120 with neMESYS pumps, Germany) and glass syringes (Hamilton, USA) were used to inject the aqueous and oil phases. Aqueous phases merged with oil phase at T-junction and generated droplets in a continuous mode of device operation (SI, Figure S11A). Solution of MTs/kinesin-1 and ATP module encapsulated continuously inside water-in-oil droplets at T-junction in a stabilized manner (SI, Figure S11B) and passed from red to green area of the channel where the height of the micro-channel is reduced to 17 μm to flatten the droplets which helps in microscopy (SI, Figure S11C). Further, droplets were entrapped between pillars in observation chamber by applying the pressure at top layer which aided in blocking the forward flow (SI, Figure S11D). Inside observation chamber entrapped-encapsulated MTs/kinesin-1/ATP module were imaged by epi-fluorescence microscope to observe contraction of filamentous network. The size of droplets was measured to be 50 ± 20 μm in diameter.

### Fabrication of the microfluidic device

A schematic presentation of the microfluidic device is shown in Figure S11A. Polydimethylsiloxane (PDMS) microchannel was fabricated using conventional soft lithography. A thin layer of SU-8 3010 photoresists (MicroChem, Newton, MA) was spin-coated onto a Si wafer and patterned via ultraviolet exposure through a chrome mask. The device has two different heights (red and green parts in Figure S11A), thereby spin-coating is done in two steps. To fabricate the device, PDMS and the curing agent (SYLGARD 184 Silicone Elastomer) were thoroughly mixed with 10:1 and poured onto the patterned SU-8 mold, followed by degassing to remove bubbles from the PDMS under vacuum pressure. After removing the bubbles and baking at 75°C for 45 min, PDMS was peeled off from the Si substrate, and the microchannels were cut to the appropriate sizes. After punching the inlet and outlet ports, PDMS microchannel was oxygen plasma-bonded (PDC 002, Harrick Plasma, Ithaca, USA) to the glass slide (24 mm×60 mm, Menzel gläser, Germany) for 30s at 200 W and 200 mTorr. To strengthen the bonding, the device was heated in a 75°C oven for at least 2 min. The height of the observation chamber is 13 μm, respectively, whereas the width was 120 μm. Diameter of pillars and pillar to pillar distance inside observation chamber was 30 and 150 μm, respectively. Micro-channel was coated with Novec^TM^ 1720, 3M at 120 °C for 1 hr to make it hydrophobic.

### Light-triggered glucose consumption

For measurement of light-triggered glucose consumption a solution containing the ATP module was supplemented with ADP, P_i_, glucose and hexokinase. As a control, a solution containing only the ATP module, ADP and Pi was run in parallel. The reaction was started by illumination and samples were taken each 5 minutes from the reaction mixture. The reaction was stopped by the addition of trichloracetic acid. The ATP concentration of the sample and the control was calculated and the reaction rates were determined by linear regression (Figure S12A). The suspension containing the ATP module, hexokinase and glucose (ATP module+hexokinase+glucose) showed barely any ATP production (0.01 nM/min). All ATP is constantly consumed by hexokinase to convert glucose in glucose-6-phosphate. This result confirmed that the concentration of hexokinase was chosen high enough and consequentially the glucose consumption reaction was not the limiting step. In contrast, the control sample (ATP module) verified that the ATP module was working and that it produced 58.6 nM ATP per min. The glucose concentration was determined using a highly sensitive glucose assay, which allows the detection of glucose by measuring the fluorescence intensity. Therefore, a standard curve relating glucose concentration and fluorescence signal was taken according to the manufacturers protocol. The glucose concentration over time is shown in Figure S12C. The glucose concentration at *t* = 0 complies well with the concentration of glucose added to the solution (10μM), which certifies that the determination of glucose with the highly sensitive assay worked. The concentration of glucose was decreasing over time with a consumption rate of −69.4 nM glucose per min. This rate is slightly higher (18%) compared to the detected ATP production rate. This result is as expected because ATP synthase is product inhibited by ATP itself^85^. Thus, higher turnover rates are expected when ATP is constantly removed from the system.

## Acknowledgment

R.A., V. N., E. B., I. G. and A. G. acknowledge support from the European Union’s Horizon 2020 research and innovation programme under grant agreement MAMI No. 766007. C.K., A.B., E.B., K.S., I.G., T.V.K., and A.G. thank MaxSynBio Consortium, which is jointly funded by the Federal Ministry of Education and Research of Germany and the Max Planck Society. We would like to thank the unknown referees, and Anthony Vecchiarelli (Assistant Professor, University of Michigan) and class MCDB 401 (“Building the Synthetic Cell”) for conducting a class review of our preprint paper, providing us with constructive and encouraging feedback. A. G. and R. A. also thank M. Lorenz and S. Bank and the Göttingen Algae Culture Collection (SAG) for providing the *Chlamydomonas reinhardtii* strain 11-32b.

## Author contributions

A.G. and T.V.K. designed the project. R.A. and A.G. isolated flagella and C.K., and T.V.K. built the energy module. R.A. and C.K. integrated the energy and flagella motility unit. R. A., V. N. and I. G. performed experiments with microtubules. R.A., C.K., Y.S., S.G.P. and A.G. analyzed the flagella data. A. G. performed the mode analysis of axonemes and A.B. wrote the Matlab code to track the axonemes and designed the microfluidic set up for droplet generation. R.A., C.K., V. N., I. G., T.V.K. and A.G. wrote the first draft of the manuscript and all the authors discussed the results and contributed to the revised version.

## Supplementary Information

**Figure S1:**
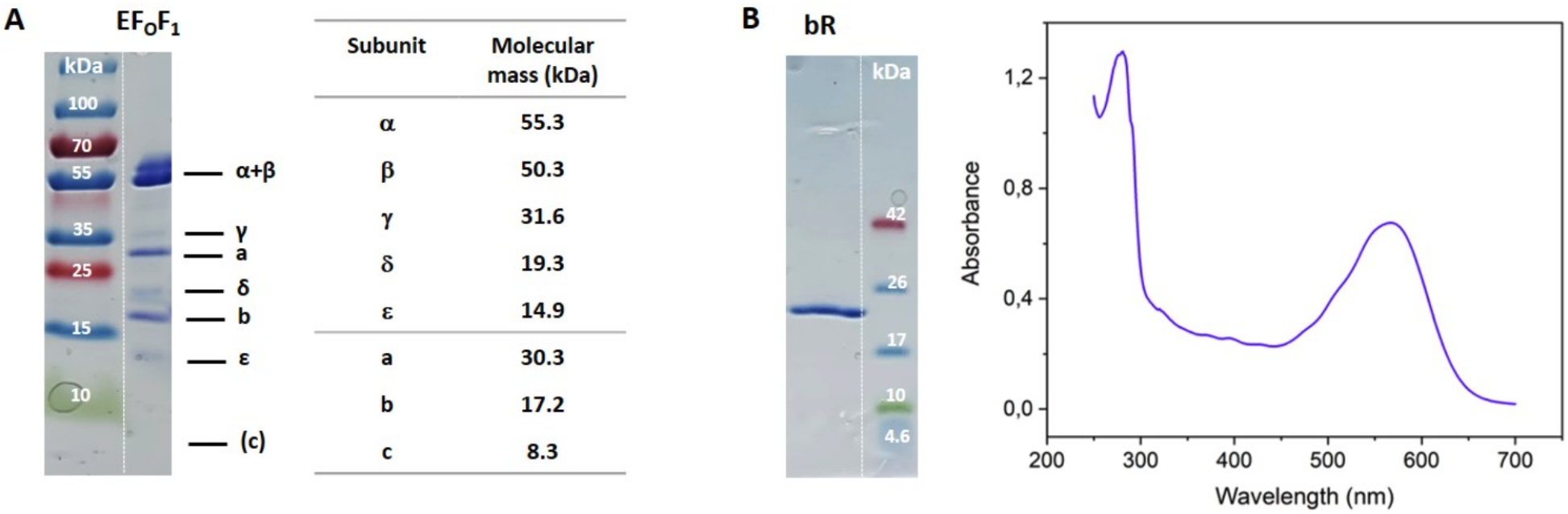
SDS-PAGE of purified proteins. (A) Coomassie-stained SDS-PAGE of purified EF_O_F_1_-ATP synthase and molecular masses of EF_O_F_1_ subunits. (B) Coomassie-stained SDS-PAGE of purified bR with the corresponding absorbance spectrum.

**Figure S2:**
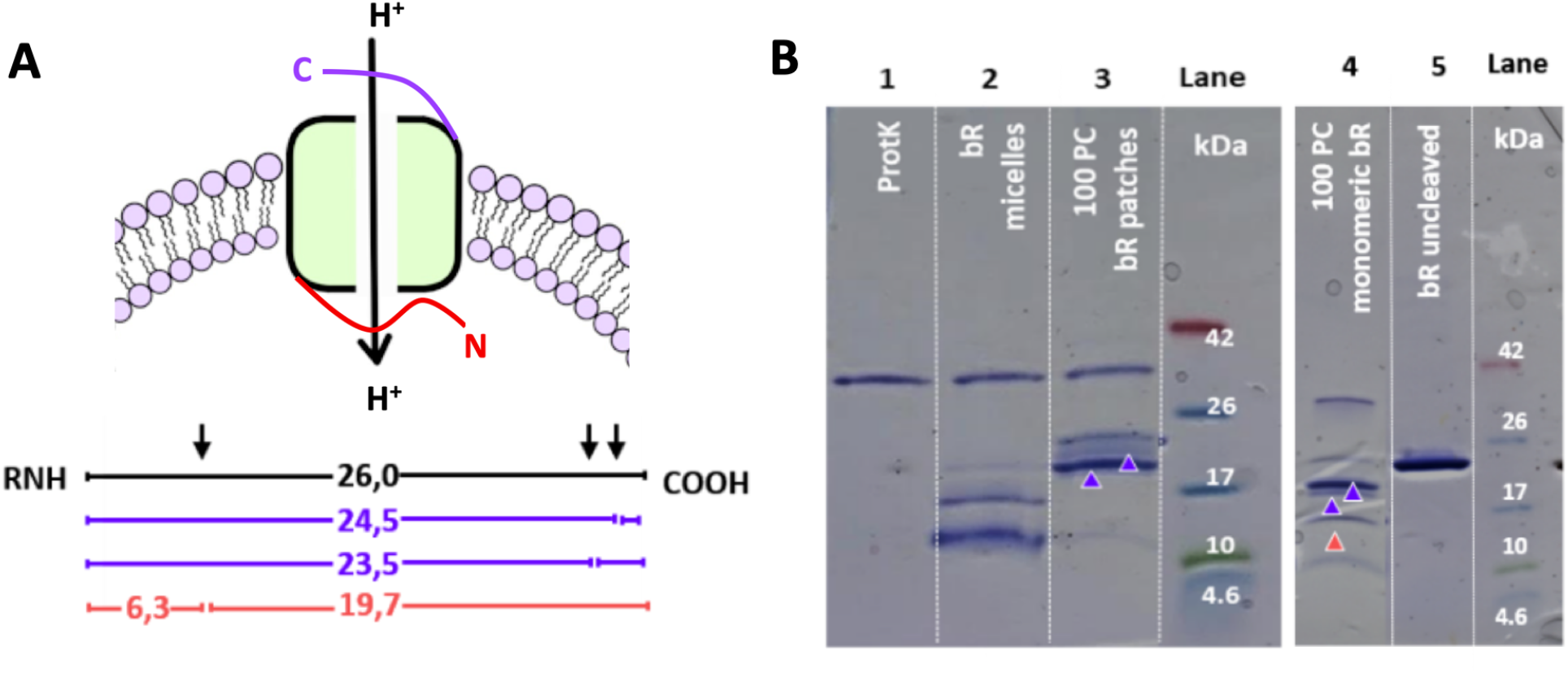
Proteolytic cleavage of reconstituted bR with proteinase K (ProtK) shows mixed orientation for monomeric bR and almost uniform orientation (with the c terminus outwards) for bR patches. (A) Expected sizes of proteolytic fragments for ProtK digestion of bR when the N-terminal (orange values) or C-terminal (violet values) is exposed to the bulk solution (B) SDS-PAGE gel analysis of the digestion products. Lane 1: band specific for ProtK enzyme only; lane 2: digestion product of not reconstituted bR; lane 3: digest pattern for PC vesicles containing bR patches; lane 4: digest pattern for PC vesicles containing monomeric bR; lane 5: undigested bR in PC lipid vesicles. Part of this data was previously published in Ref.^36^.

**Figure S3:**
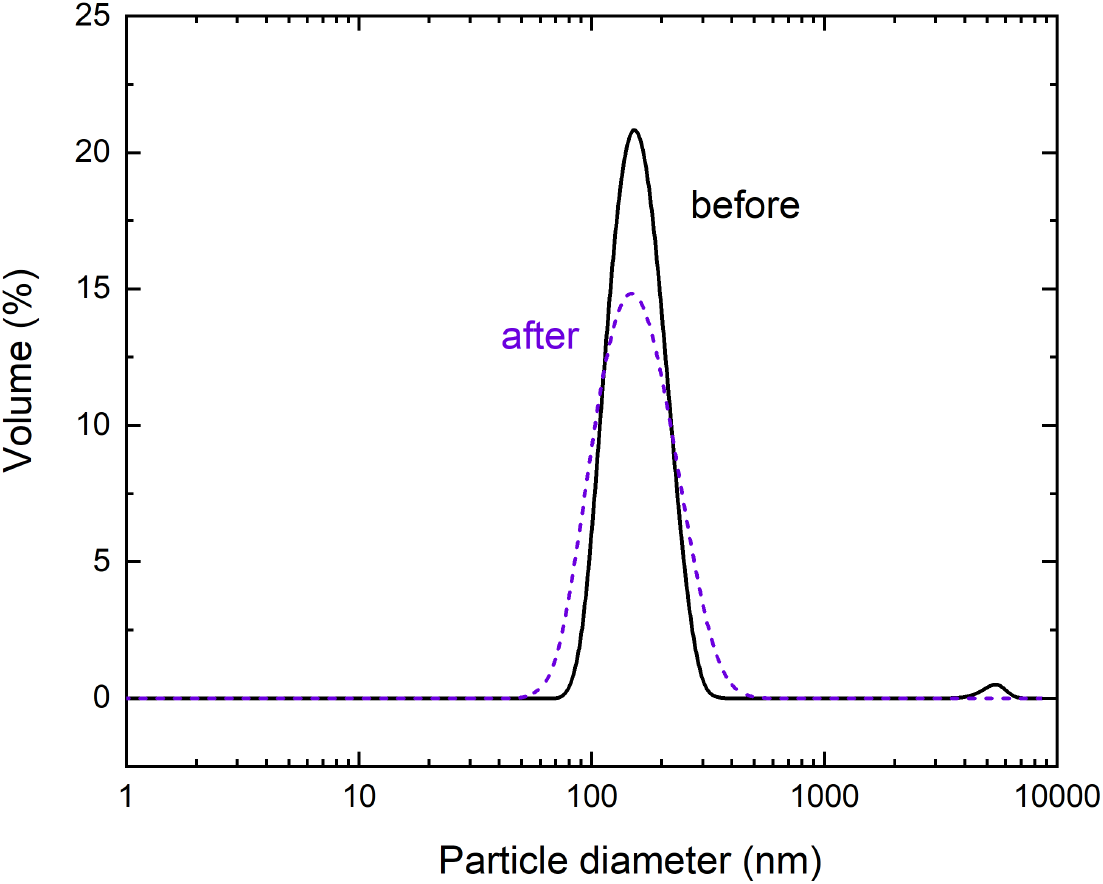
Size distribution of vesicles before (before) and after reconstitution and removal of detergent using Bio Beads (after).

**Figure S4:**
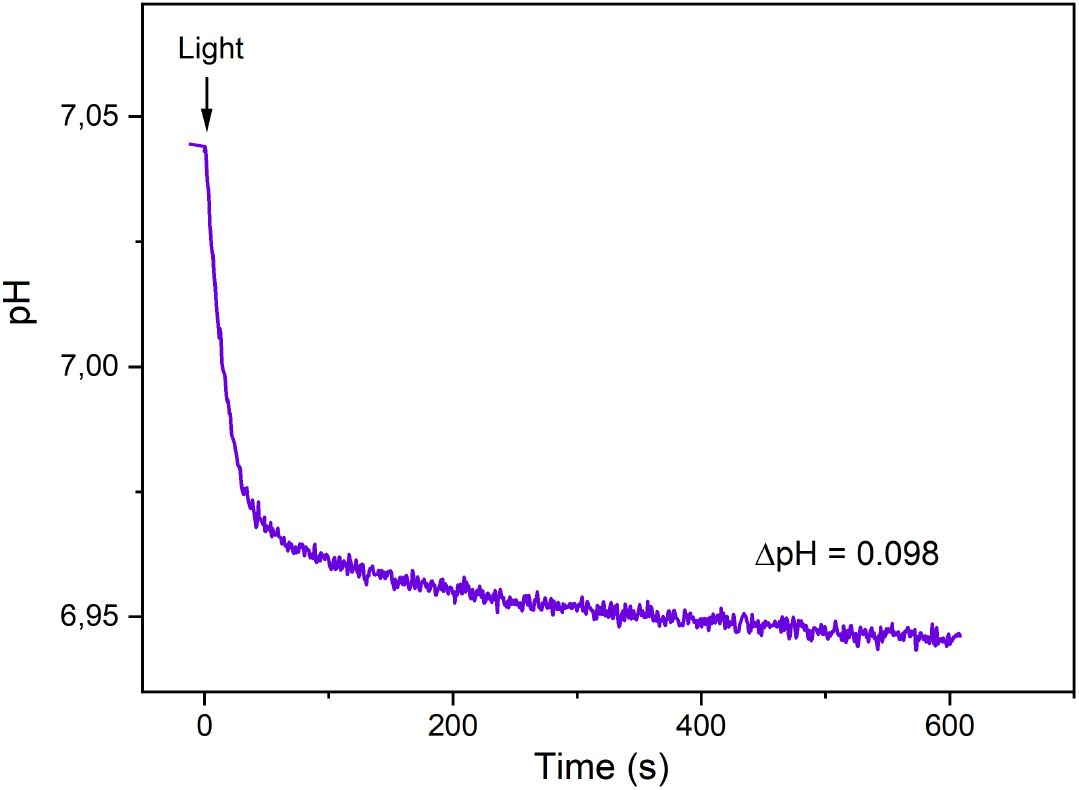
Proton pumping of bR as measured by encapsulated pyranine.

**Figure S5:**
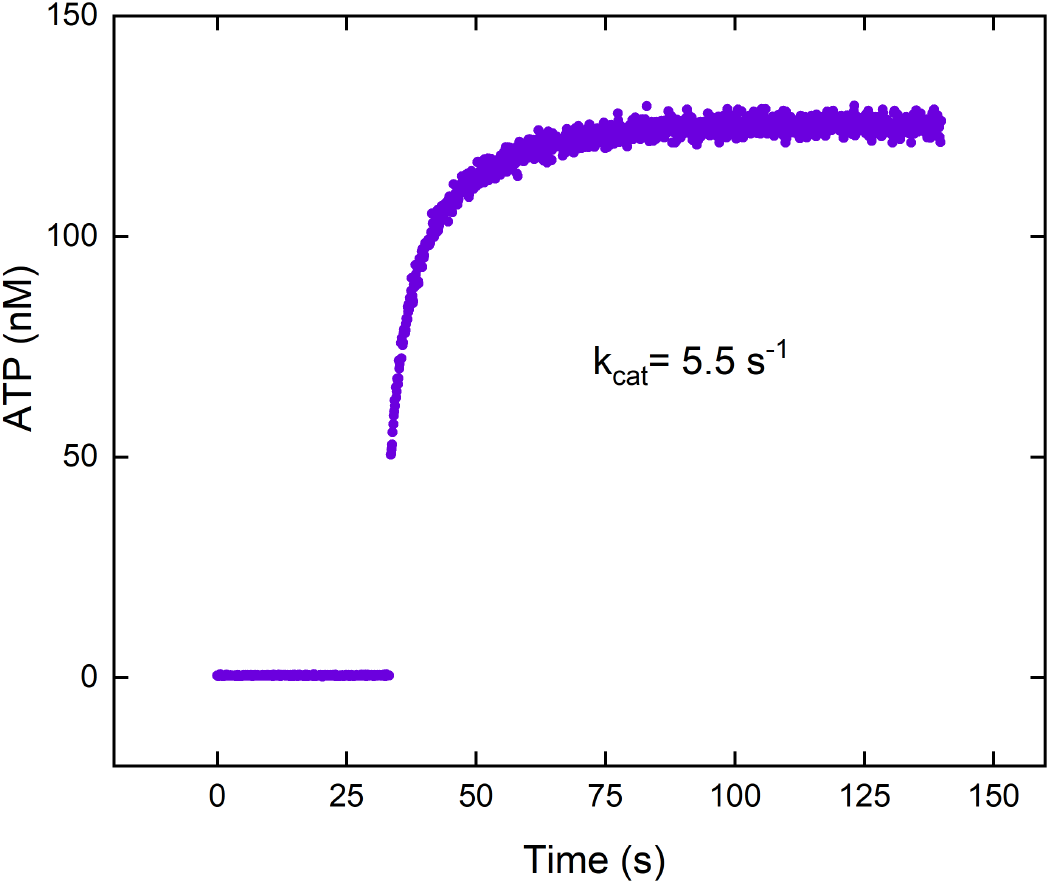
Turnover (k_*cat*_) of ATP synthase as determined in an acid-base transition experiment. Measurements were performed in 20 mM succinate-NaOH, 5 mM NaH_2_PO_4_, 0.6 mM KOH (inner solution) and 200 mM tricine-NaOH, 5 mM NaH_2_PO_4_, 160 mM KOH (outer solution) adjusted with 18 μM valinomycin at room temperature. [ADP]= 0.1 μM, [P^i^]= 5 mM, [lipid]= 0.24 mg/mL, [EF_0_F_1_]= 2.22 nM, ΔΨ= 143 mV. ATP synthase was reconstituted with 0.8 % Triton.

**Figure S6:**
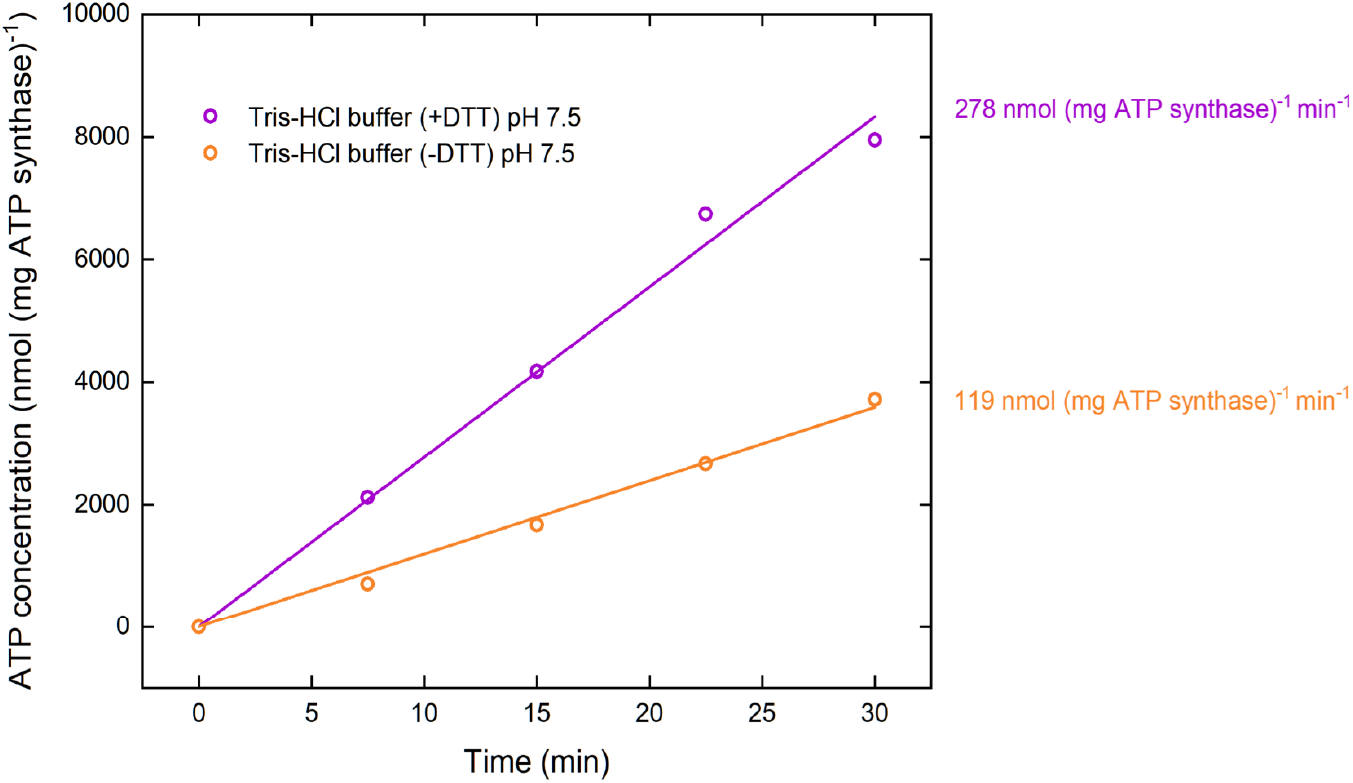
ATP production in Tris-HCl buffer in the presence (+DTT) and absence of DTT (-DTT). DTT conserves proteins in their functional form as it prevents oxidation of sulfhydryl groups (SH-) to disulfide-bonds in the presence of air oxygen. The activity was determined by linear regression. Measurements were performed with [ADP]= 300 μM, [P_i_]= 5 mM, [lipid]= 0.022 mg/mL, [EF_0_F_ļ_]= 1.3 nM, [bR]= 88 nM, ΔΨ= 143 mV at room temperature. Proteins were reconstituted with 0.8 % Triton.

**Figure S7:**
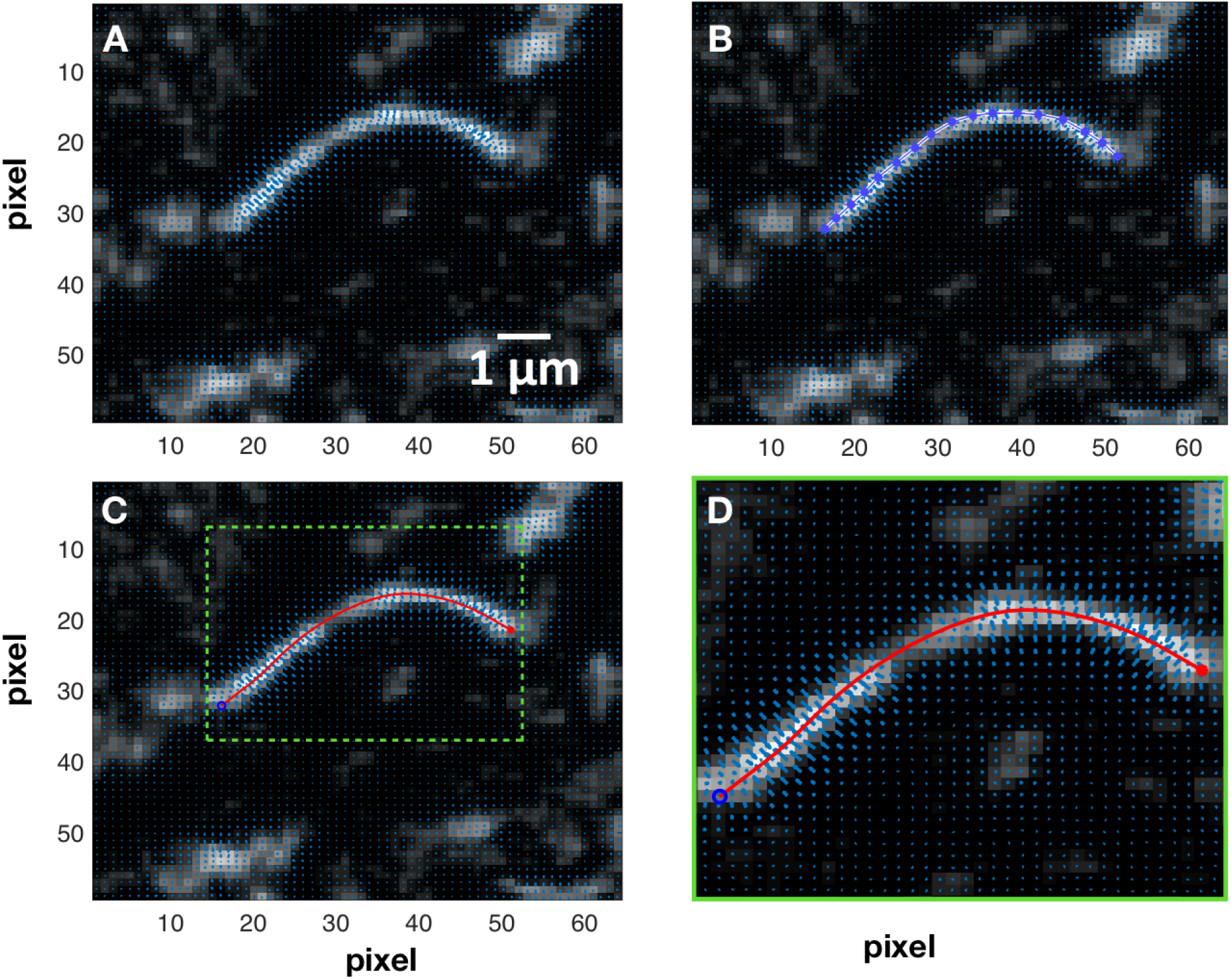
A) Gradient vector flow calculated at the vicinity of an axoneme. B) The initial selection of a polygon for the first frame which deforms according to the gradient vector flow. C) The final tracked shape of an axoneme. D) A zoomed-in image showing the gradient vector flow (blue arrows) at a higher magnification.

**Figure S8:**
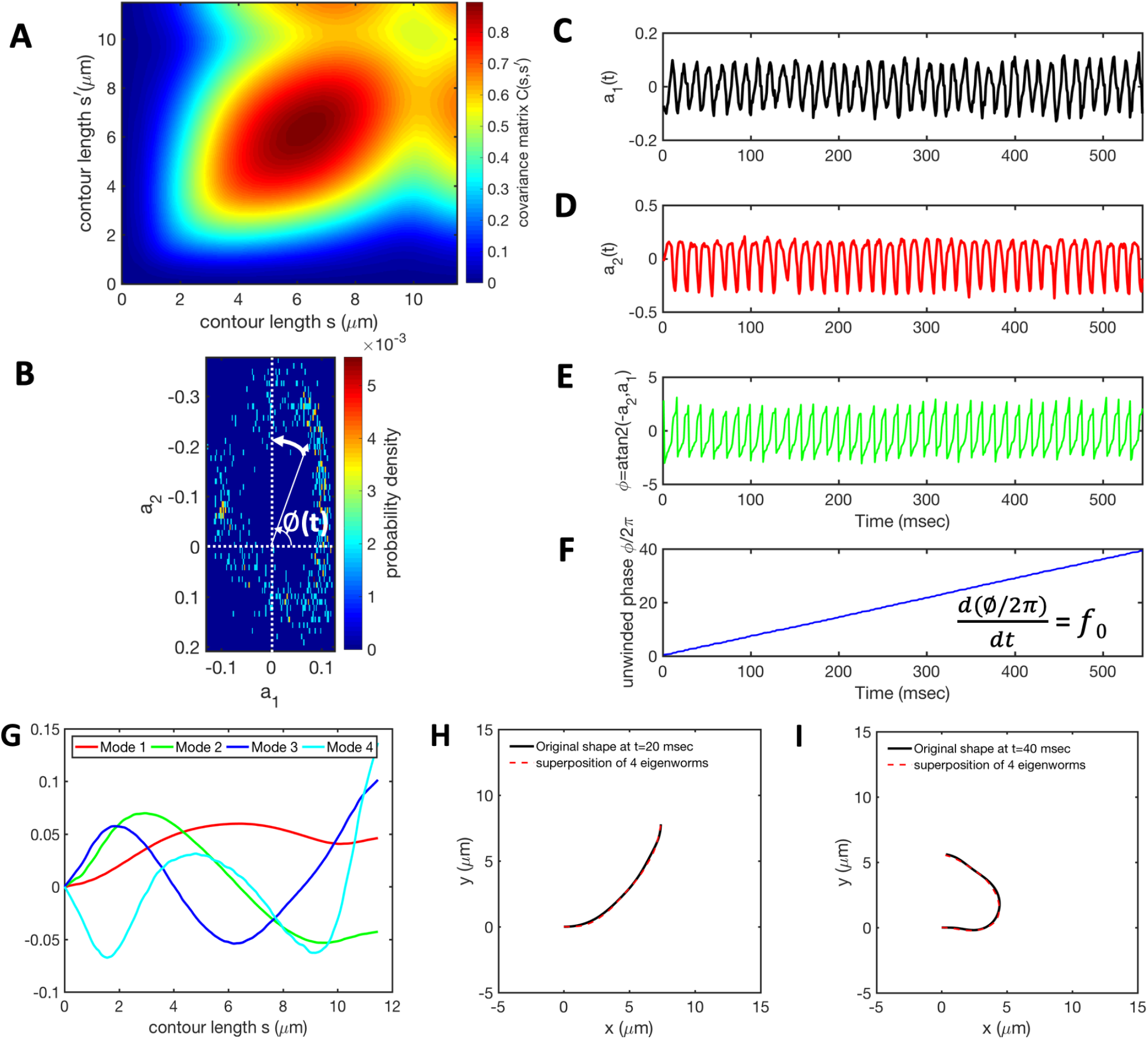
Mode analysis of the reactivated axoneme presented in Figure 6A-E. A) The covariance matrix *C*(*s, s*’) of fluctuations in angle *θ*(*s,t*). B) The probability distribution of the first two shape amplitudes *p*(*a*_1_(*t*), *a*_2_(*t*)). The phase angle of the axoneme as an oscillator is defined as *ϕ*(*t*) = atan2(-*a*_2_(*t*), *a*_1_(*t*)). C-D) Time evolution of the first two dominant shape amplitudes *a*_1_(*t*) and *a*_2_(*t*) showing regular oscillations at frequency of 72 Hz. E-F) We observe a linear growth in dynamics of *ϕ*(*t*), indicating steady rotation in *a*_1_ — *a*_2_ plane presented in part B. Note that *dϕ/dt* = 2*πf*_0_ where *f*_0_ is the beating frequency of axoneme. G) Four eigenvectors corresponding to the four largest eigenvalues of matrix *C*(*s, s*’). H-I) Superposition of four eigenmodes presented in part G with coefficients *a*_1_(*t*) to *a*_4_(*t*), can reproduce shape of the axoneme with high accuracy.

**Figure S9:**
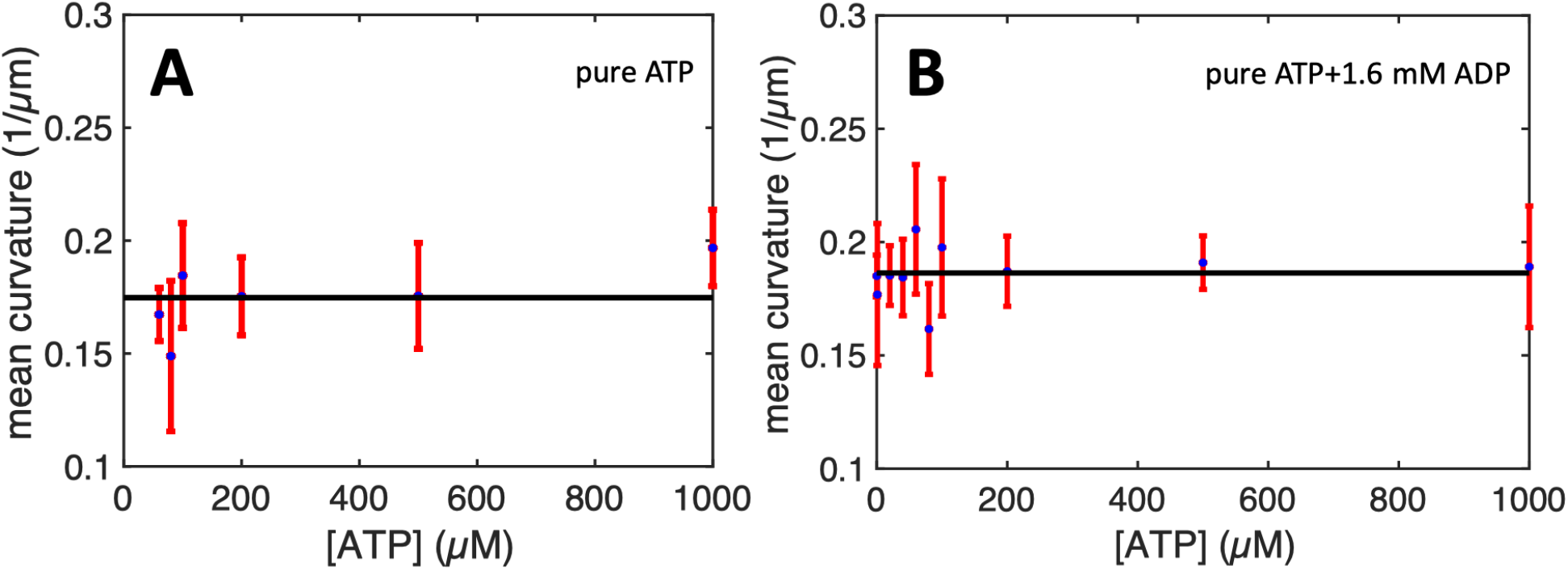
Mean curvature of axonemes reactivated using pure commercial ATP plotted versus ATP concentration. In panel A, no ADP is added, while in panel B, the reactivation buffer is supplemented with 1.6 mM ADP. The black solid lines show the mean values of 0.17 *μ*m^-1^ (A) and 0.186 *μ*m^-1^ (B), which is comparable to the value of 0.16 *μ*m^-1^ in energy module experiments shown in Fig. 4E. The corresponding frequency trends in pure ATP experiments with and without ADP are shown in Fig. 5C and Fig. 3E, respectively.

**Figure S10:**
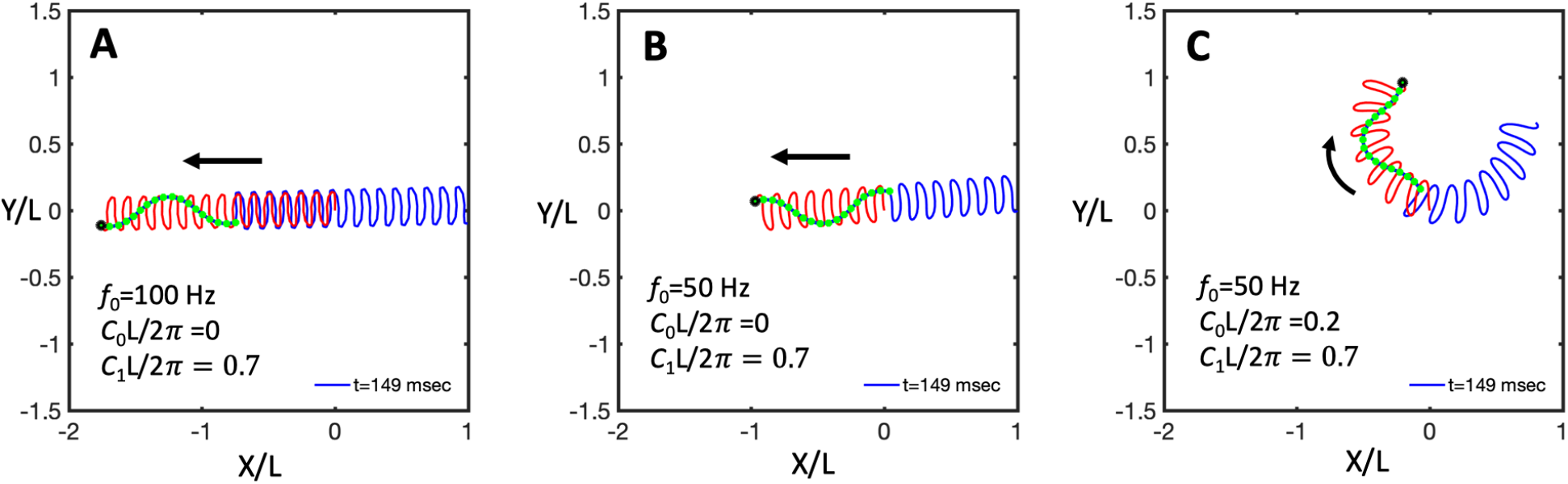
A-B) A flagellum swimming in the absence of static curvature *C*_0_ at two different frequencies of 100 and 50 Hz. Faster beating flagellum swims a longer distance. C) Flagellum follows a circular path if *C*_0_ is non-zero. Amplitude of the dynamic mode *C*_1_ is kept the same for all three simulations. *L* is the contour length of flagellum (see Video 8).

**Figure S11:**
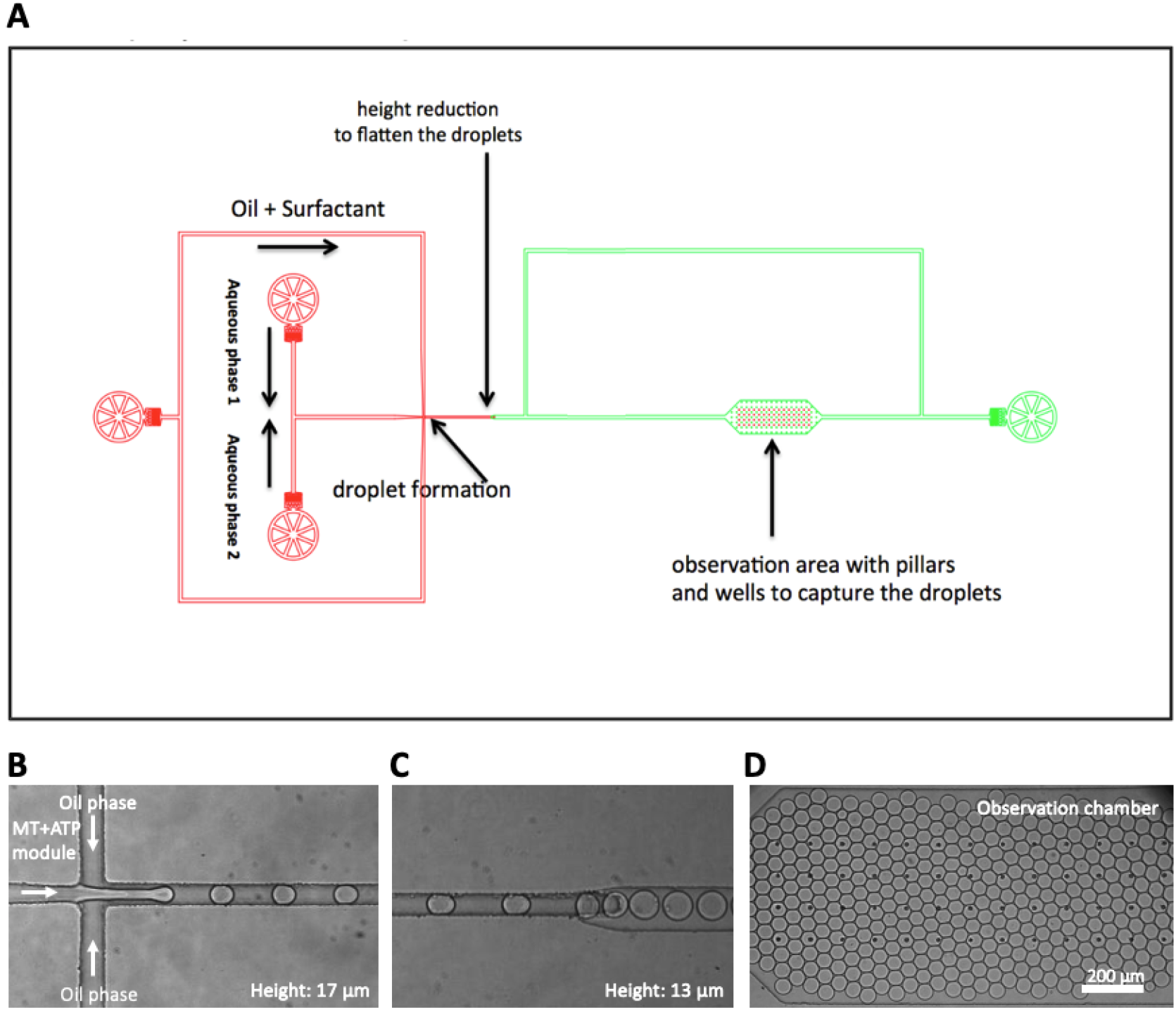
Droplet microfluidic setup. A) Schematic diagram showing the encapsulation of MTs/kinesin-1 network (aqueous phase 1) with light-driven ATP module (aqueous phase 2). B) Mono-disperse water-in-oil droplets containing MTs/kinesin-1 solution and light-driven ATP module are formed at the T-junction. C) Droplets enter the green area where height is decreased from 17 *μ*m to 13 *μ*m. D) Observation chamber with entrapped droplets where the flow into the chamber was stopped by guiding the flow to the side channel. The following flow rates were adjusted: aqueous phase 1, 200 *μ*L/h; aqueous phase 2, 200 *μ*L/h; oil phase, 400 *μ*L/h.

**Figure S12:**
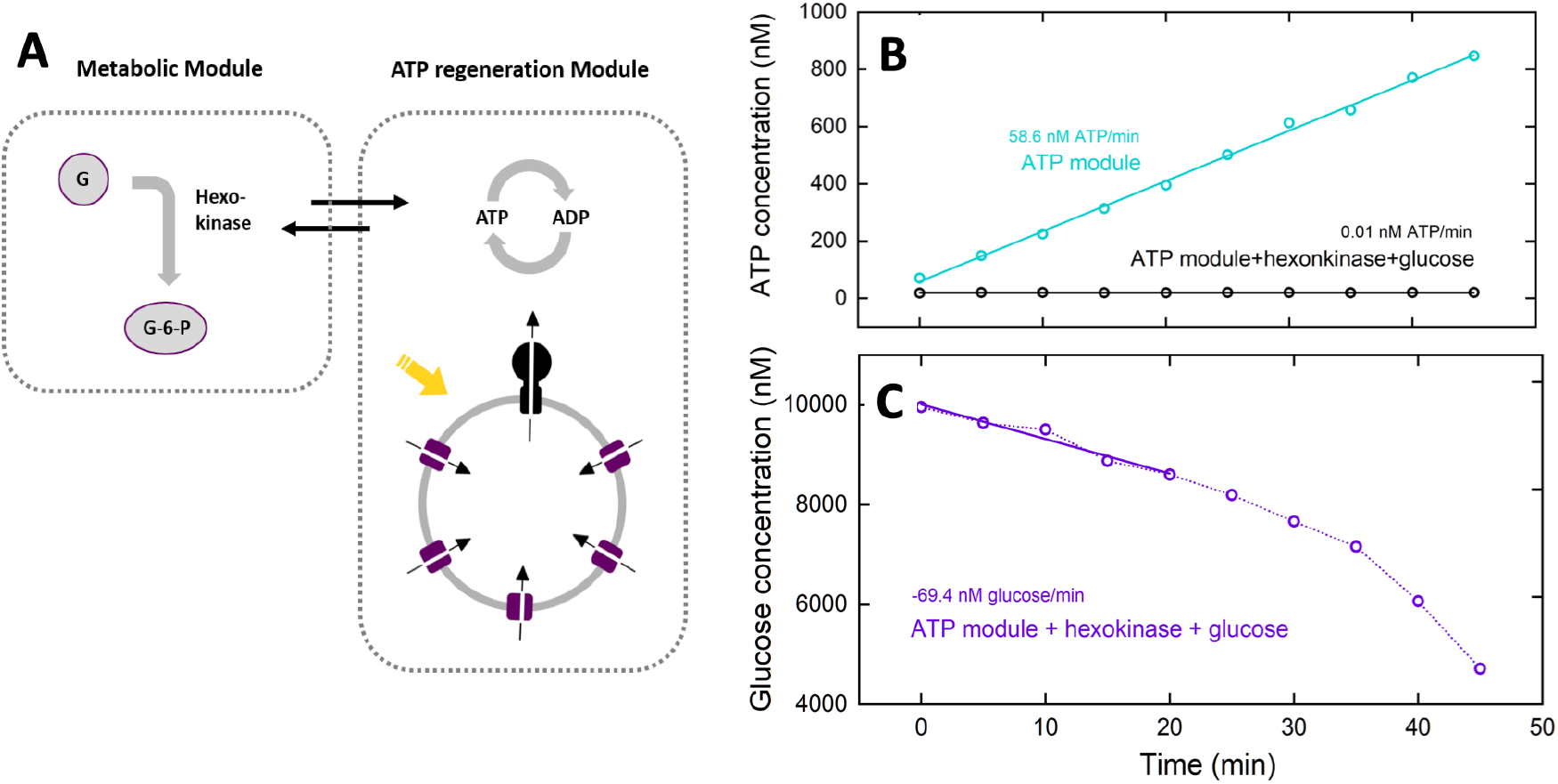
ATP and glucose concentration over time under green light illumination. A) Scheme of coupling the ATP regeneration module with a metabolic module for glucose consumption triggered by light. B-C) Measurements were performed in 20 mM tricine-NaOH, 20 mM succinate, 0.6 mM KCl, 80 mM NaCl (inner solution) and 200 mM tricine-NaOH, 5 mM NaH_2_PO_4_, 160 mM KOH, pH 8.8 (outer solution) in the presence of 20 μM valinomycin at room temperature. [lipid]= 0.022 mg/mL, [EF_O_F_1_]= 1.3 nM, [bR]= 88 nM, ΔΨ= 143 mV. Proteins were reconstituted with 0.8 % Triton. For measurement of glucose consumption 10 μM glucose and 2.7 μg/mL hexokinase (2400 U/mL) was added to the outer solution. Glucose concentration was determined using a highly sensitive glucose assay (Sigma).

**Figure S13:**
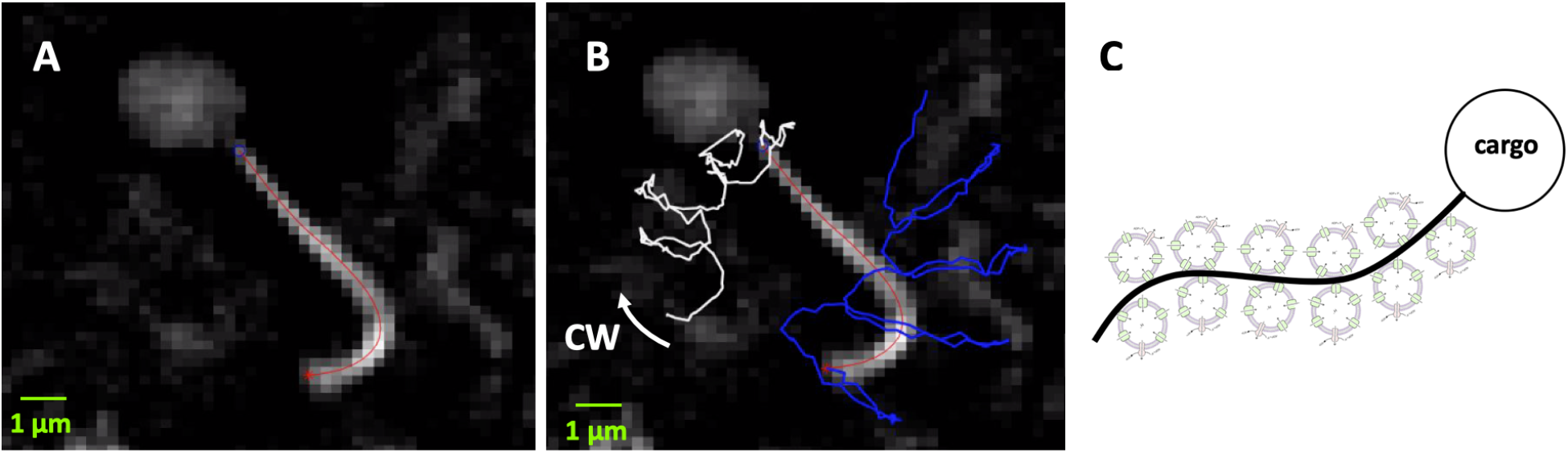
Bead as a cargo. A) A bead of radius 1 *μ*m is attached to the distal end of an axoneme. B) Upon illuminations, the reactivated axoneme propels the bead. Blue line shows the trace of the basal end and the white line is the trace of the distal tip which is attached to a bead (SI, Video 11). C) A cartoon showing that an enhanced vesicle-axoneme attachment, e.g. via electrostatic interactions, could be beneficial for the local production of ATP in the vicinity of axonemes.

**Figure S14:**
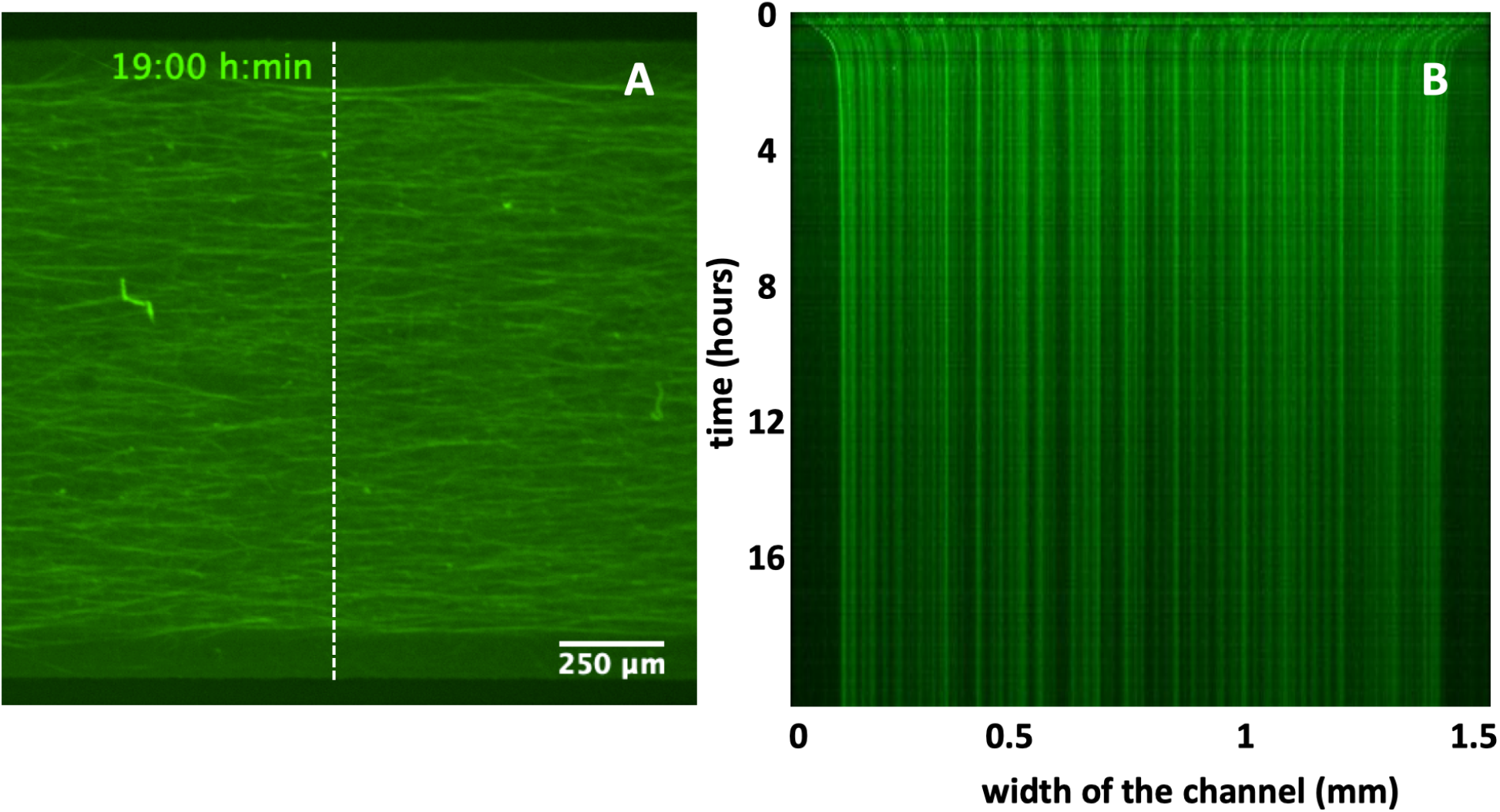
A control experiment with 1 mM pure commercial ATP to confirm that a contracted MTs/kinesin1 network does not relax back to its initial state once the ATP is depleted. A) Snapshot of the millifluidic set up filled with MTs/kinesin1 network after 19 hours. B) The space-time plot along the white dashed line in panel B showing the network contraction. Note that the cross-linked network remains contracted for a long period of time.

